# Butyrate Rescues Cardiac Metabolic Dysfunction in Hypertensive Heart Failure with Preserved Ejection Fraction

**DOI:** 10.1101/2025.11.28.690152

**Authors:** Sarah M. Kedziora, Sabrina Y. Geisberger, Oliver Popp, Alina Frovola, Alisa Uhrbach, Kristin Kräker, Nurana Tagiyeva, Marcus Kelm, Nikolaus Berndt, Yiming D. Zhang, Guido Mastrobuoni, Raquel Escrihuela-Branz, Theda U. P. Bartolomaeus, Lorenz Pixner, Pauline Fahjen, Gelsomina N. Kaufhold, Pragati Parakkat, Alexandra M. Chitroceanu, Carsten Tschöpe, Frank Edelmann, Gabriele G. Schiattarella, Sofia K. Forslund, Nicola Wilck, Michael Gotthardt, Philipp Mertins, Ralf Dechend, Suphansa Sawamiphak, Dominik N. Müller, Stefan Kempa, Nadine Haase

**Affiliations:** Max Delbrück Center for Molecular Medicine in the Helmholtz Association, 13125 Berlin, Germany; Experimental and Clinical Research Center, a cooperation of Charité-Universitätsmedizin Berlin and Max Delbrück Center for Molecular Medicine, 13125 Berlin, Germany; Charité - Universitätsmedizin Berlin, corporate member of Freie Universität Berlin and Humboldt-Universität zu Berlin, 10117 Berlin, Germany; DZHK (German Centre for Cardiovascular Research), partner site Berlin, 10785 Berlin, Germany; Integrative Proteomics and Metabolomics, Berlin Institute for Medical Systems Biology, Max Delbrück Center for Molecular Medicine in the Helmholtz Association, 13125 Berlin, Germany; Proteomics Platform, Max Delbrück Center for Molecular Medicine in the Helmholtz Association, 13125 Berlin, Germany; Institute of Computer-assisted Cardiovascular Medicine, Deutsches Herzzentrum der Charité, 13353 Berlin, Germany; Department of Congenital Heart Disease – Pediatric Cardiology, Deutsches Herzzentrum der Charité, 13353 Berlin, Germany; Department of Molecular Toxicology, German Institute of Human Nutrition Potsdam-Rehbruecke (DIfE), 14558 Nuthetal, Germany; Department of Radiology, Charité – Universitätsmedizin Berlin, corporate member of Freie Universität Berlin and Humboldt-Universität zu Berlin, 13353 Berlin, Germany; Deutsches Herzzentrum der Charité (DHZC), Department of Cardiology, Angiology and Intensive Care Medicine, 13353 Berlin, Germany; Max Rubner Center for Cardiovascular Metabolic Renal Research (MRC), Charité-Universitätsmedizin Berlin, Germany, 10115 Berlin, Germany; Friede Springer Cardiovascular Prevention Center at Charité – Universitätsmedizin Berlin, 12203 Berlin, Germany; Division of Cardiology, Department of Advanced Biomedical Sciences, Federico II University, Naples, Italy; Charité - Universitätsmedizin Berlin, Department of Nephrology and Internal Intensive Care Medicine, 13353 Berlin, Germany; Core Unit Proteomics, Berlin Institute of Health at Charite - Universitätsmedizin Berlin, 10117 Berlin, Germany; HELIOS Clinic, Department of Cardiology and Nephrology, 13125 Berlin, Germany

## Abstract

Diastolic dysfunction in heart failure with preserved ejection fraction (HFpEF) is characterized by metabolic inflexibility. Unlike systolic heart failure, where ketone bodies support energy homeostasis, the failing heart in HFpEF lacks well-characterized alternative fuels to meet its high ATP demand. Here, we show that butyrate, a microbiota-derived short-chain fatty acid, serves as an ancillary energy source and improves diastolic function. Although cardiac power was preserved in rats with HFpEF, both experimental and human HFpEF hearts exhibited an impaired expression of proteins in mitochondrial electron transport chain and oxidative phosphorylation. Additionally, accumulation of 3-hydroxy-butyrate (BOH) in rat and also human HFpEF indicated that ketones do not rescue the cardiac energetic deficit. In HFpEF patients from the UK Biobank, higher BOH levels were associated with increased mortality, particularly those with hypertension. Applying ^13^C-butyrate to isolated perfused hearts with and without HFpEF resulted in isotope incorporation in butyryl-CoA and downstream TCA intermediates and thus proving its active metabolization. Butyrate was efficiently oxidized by cardiomyocytes and was overtaking BOH and amino acids in supporting respiration. Finaly, chronic butyrate supplementation improved survival, enhanced diastolic function, and reduced fibrosis and inflammation in HFpEF rats despite persistent hypertension. These findings identify butyrate as a compensatory fuel and a promising therapeutic candidate in energetically compromised HFpEF.

## Main

Life depends on energy. Mitochondria, often referred to as the powerhouses of the cell, play a pivotal role in both health and disease. These organelles are present in nearly all human cells and generate ATP through oxidative phosphorylation. They originated approximately 1.5 to 2 billion years ago through endosymbiosis between an aerobic α-proteobacterium and an archaeal host, establishing the foundation for eukaryotic energy metabolism. In 1861, Pasteur discovered butyrate as a fermentation product of obligate anaerobes, while its importance as a primary energy source for colonocytes only became clear in the late 20th century. However, its role as an energy substrate in the heart remains poorly understood.

Heart failure (HF) with preserved ejection fraction (HFpEF) is a growing health burden linked to aging, obesity, diabetes, and hypertension^1^ and now accounts for over half of all HF hospitalizations. In contrast to heart failure with reduced ejection fraction (HFrEF), availability of therapeutic options for HFpEF is limited, reflecting its multifactorial and incompletely understood pathophysiology. Energetic impairment is a hallmark of the failing myocardium, but metabolic adaptations in HFpEF have not been fully elucidated. Cardiac contraction is one of the most energy-intensive processes in the body, requiring more than 6 kg of ATP per day for lifelong contraction^2^. In the healthy heart, over 90% of this ATP is generated by mitochondrial oxidative phosphorylation with fatty acids serving as the predominant fuel source^3^. While glucose is more oxygen efficient per ATP, this shift does not sufficiently meet the demand. HFrEF hearts partially compensate by increasing reliance on alternative fuels such as ketone bodies^1,4^, but whether similar adaptations occur in HFpEF and if butyrate has a therapeutic potential *in vivo* remained unclear.

Beyond the traditional heart-centric view, there is growing evidence that the gut microbiome is a key regulator of interorgan communication. Through fermentation of dietary fiber, commensal bacteria produce SCFAs, including acetate, propionate, and butyrate, which function as locally and systemically active metabolites^5^. SCFAs exert broad effects on host metabolism, inflammation, endothelial function, and immune regulation, positioning them as critical intermediaries between diet, microbial ecology, and cardiometabolic disease^5^. Disruptions in the composition or function of the gut microbiota have been linked to cardiovascular pathologies, including hypertension, obesity, and HF^6,7^. Patients with HFpEF show reduced levels of SCFA-producing microbes^8^ and lower circulating SCFA concentrations^9^. In humans, prebiotic fiber interventions that increase SCFA production lower blood pressure^10^, a core risk factor for HFpEF. Low SCFA levels are associated with increased intestinal permeability, inflammation^11^, and endothelial dysfunction^12^, all pathophysiological features of HFpEF. In rodents, butyrate improves myocardial energetics in high sucrose-treated mice^13^, attenuates fibroblast differentiation^14^ and cardiac fibrosis by shifting macrophage polarization^14^. Although failing HFrEF myocardium can efficiently oxidize butyrate, surpassing even ketone bodies as a preferred substrate^15,16^ - its role in HFpEF remains unresolved.

Here, we address this gap using a hypertensive HFpEF rat model in combination with proteomics, metabolomics, ^13^C-butyrate tracing, and kinetic metabolic modeling. We hypothesize that butyrate serves as an alternative fuel that mitigates myocardial energy deficiency and improves diastolic function in HFpEF. Our findings support butyrate supplementation as a mechanistically targeted metabolic strategy for treating HFpEF.

### Metabolic rewiring of the hypertensive HFpEF hearts

The progression of HFpEF is characterized by a gradual deterioration of cardiac structure and function. To model this trajectory, we used double transgenic rats (dTGR) overexpressing both human renin and angiotensinogen, thereby mimicking a severe progressive form of hypertensive diastolic dysfunction^17^. Untreated dTGR exhibited markedly reduced survival by seven weeks of age and progressive weight loss compared to healthy, non-transgenic controls (WT; Extended Data Fig. 1a and 1b), reflecting end-stage HFpEF^17–22^. In untreated dTGR, left ventricular EF was preserved (Fig. 1a) despite steadily increased systolic blood pressure (BP) (Fig. 1b). Echocardiography revealed pronounced concentric remodeling with posterior wall thickening (Fig. 1c) and increased interstitial fibrosis (Extended Data Fig. 1c), consistent with maladaptive structural remodeling characteristic of human hypertensive HFpEF, in which chronic pressure overload drives concentric geometry and extracellular matrix expansion^23^. Invasive hemodynamic assessment of cardiac performance demonstrated leftward-shifted pressure-volume loops (Fig. 1d), reduced stroke volume (Fig. 1e), and elevated arterial elastance and total peripheral resistance (Fig. 1f, 1g), reflecting diastolic dysfunction comparable to that observed in patients with hypertensive HFpEF. Global longitudinal strain (GLS), a sensitive marker of HFpEF^24^ was significantly impaired in dTGR (Fig. 1h), with the most pronounced regional dysfunction localized to the posterior apex (Fig. 1i).

**Fig. 1.**
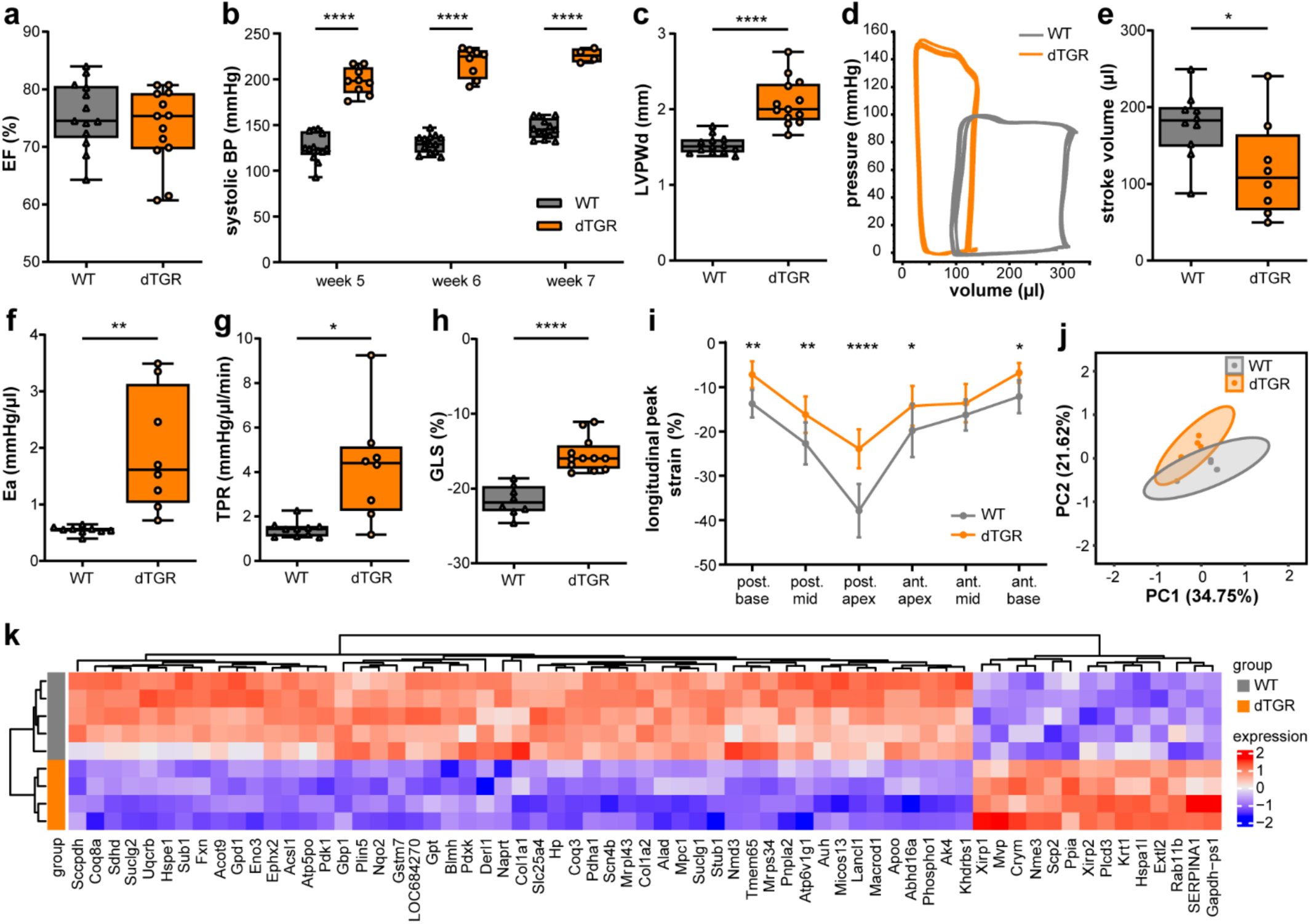
Hypertensive diastolic dysfunction, hypertrophy, and region-specific proteomic remodeling in a HFpEF model. **(a)** Preserved ejection fraction (EF) measured by echocardiography (n=13 each). **(b)** Elevated systolic blood pressure measured by tail-cuff at five, six and seven weeks of age. (n=13 WT, n=4-9 dTGR) **(c)** Cardiac hypertrophy assessed in echocardiography through LV posterior wall thickness in diastole (LVPWd, n=13 each), **(d)** Representative LV ventricular pressure-volume loops measured with the Millar Tip catheter system are displayed for one animal per group. **(e)** Stroke volume (n=8-10), **(f)** arterial elastance (Ea, n=8-9) and **(g)** total peripheral resistance (TPR, n=8-10) are significantly elevated in dTGR. **(h)** Mean global longitudinal strain (GLS) was assessed in parasternal long axis (PLAX) B-mode echocardiography (n=8-12). **(i)** Segmental longitudinal peak strain in PLAX was significantly increased in dTGR in all cardiac posterior (post.) and anterior (ant.) segments (n=8-12). **(j)** Principal component analysis (PCA) of all proteins with percentage of variance explained of posterior apex based on data filtered for ≥75% valid values and subsequently imputed. Distinct separation of WT and dTGR. WT vehicle, grey; dTGR, orange; (n=4-5). **(k)** Heatmap displays unsupervised hierarchical clustering of significantly dysregulated (adj.P<0.05, |logFC|>0.5) posterior apex proteins in WT | dTGR comparison. The expression data is scaled using z-scores for better visualization. (n=4-5). Outliers were removed upon statistical testing. Data are presented as boxplots (IQR) with whiskers min to max. **(a-c and e-i)** *P<0.05, **P<0.01, ***P<0.0001 **(a,c and e-h)** unpaired two-tailed Welch’s t-test, **b)** via two-way ANOVA with Sidak multiple comparison **(i)** two-way ANOVA with Dunnett‘s multiple comparison.

In line with the regional functional changes in the posterior apex, principal component analysis of regional cardiac proteome measured by label-free proteomic profiling revealed distinct group separation only in this region (Fig. 1j, Extended Data Fig. 1d–h). Posterior apical proteomic changes correlated with the region of most severe GLS impairment. Unsupervised clustering of the proteomics data and ontological analysis identified extensive alterations in structural, electrical, and mitochondrial protein networks within the posterior apex (Fig. 1k, Extended Data Fig. 1i-1m), which were less prominent in other regions. Together, these data highlight a regional cardiac proteome-function relation for the posterior apex.

To better understand how compensatory metabolic adaptation preserves cardiac power (Extended Data Fig. 2a) and cardiac function (Fig. 1d-i), we performed single-sample gene set enrichment analysis (ssGSEA) based on the proteomic data. In the posterior apex aerobic respiration, mitochondrial respiratory electron transport and electron transport chain (ETC) complex I biogenesis (Fig. 2a, Extended Data Fig 2b-g) were found downregulated. The downregulation of complex I and synthesis of ubiquinone in dTGR HFpEF hearts (Fig. 2b) is indicative for an escape of overreduction of quinone-pool that may result in ROS-induced cardiac damage in dTGR. Proteomics of endomyocardial biopsies from HFpEF patients showed reduced expression of proteins related to mitochondrial oxidative phosphorylation and fatty acid metabolism, particular in the subgroup associated with more severe obesity^25^. When comparing these changes with our proteomic data from dTGR, we found clearly overlaying changes in the respective pathways (Fig. 2c, f) providing evidence that the dTGR rat model resembles similar metabolic reprogramming as human HFpEF. Seahorse experiments in isolated primary cardiomyocytes from dTGR showed a reduced maximal respiratory capacity upon chemical uncoupling with FCCP (Fig. 2d, Extended Data Fig. 2h). *In silico* metabolic modelling using CARDIOKIN^26^ calculated a significant reduction of ATP production capacity, indicating impaired metabolic performance in dTGR (Extended Data Fig. 2i).

**Fig. 2.**
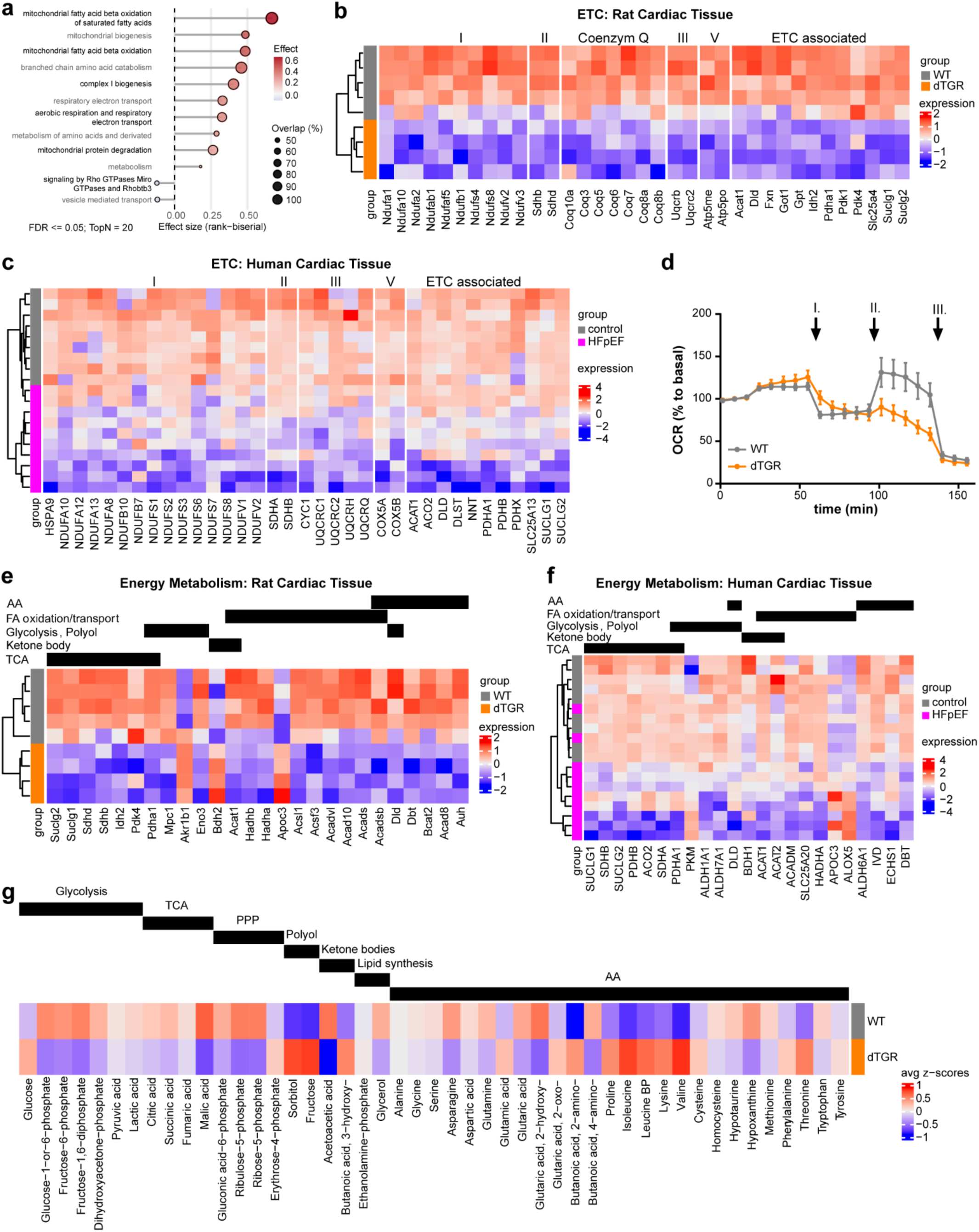
Impaired respiration and altered substrate use in dTGR hearts. **(a)** Dot plot of significantly enriched Reactome pathways based on log2 fold changes (WT vs. dTGR; n=5 WT, n=4 dTGR). Enrichment was assessed using a two-sided Wilcoxon rank-sum test with Benjamini–Hochberg correction; pathways with FDR ≤ 0.05 are shown. The x-axis indicates the rank-biserial effect size (positive = higher in WT, negative = higher in dTGR). Point size represents the percentage of quantified proteins overlapping each pathway. Pathways are ordered by the magnitude of effect size. **(b)** Hypothesis-driven heatmap of ETC-related proteins filtered by p-value ≤ 0.01 for broader biological coverage (n=5 WT, n=4 dTGR). **(c)** Heatmap of ETC-related proteins from the HFpEF human dataset filtered by p-value ≤ 0.05 (n=9 control, n=10 HFpEF). Normalized protein expression and p-values are taken from the original study. **(d)** Oxygen consumption rate (OCR; pmol/min/Hoechst intensity) measured by Seahorse technology shows comparable OCR at baseline, after injection I (oligomycin) and III (rotenone/antimycin A), with reduced OCR after injection II (FCCP) (n=30–36 technical replicates from n=5–6 experiments). **(e)** Heatmap of proteins involved in energy metabolism and substrate utilization filtered by p-value ≤ 0.01 (n=5 WT, n=4 dTGR). **(f)** Corresponding heatmap from the HFpEF dataset filtered by p-value ≤ 0.05 (n=9 control, n=10 HFpEF). **(g)** Heatmap of z-score transformed central carbon metabolites measured by GC-MS (n=3 each).

Profiling of protein expression and the central carbon metabolites in the posterior apex revealed that proteins of the TCA cycle, amino acid (AA) metabolism, fatty-acid oxidation/transport and proteins associated with oxidative phosphorylation were reduced in dTGR (Fig. 2e; Extended Data Fig. 2j-q). Intermediates of glycolysis (G6P, F6P, and F1,6P), TCA cycle, and oxidative pentose phosphate pathway were reduced in dTGR, while intracellular glucose was increased (Fig. 2g). Interestingly, despite the reduction of glycolytic intermediates, modelled glucose uptake capacity was significantly increased in dTGR in our *in silico* modelling (Extended Data Fig. 2r).

The polyol pathway, specifically aldo-keto reductase family 1 member B (AKR1B1; Fig. 2e), is one of the few increased pathways in failing dTGR apex myocardium and correlates well with increased levels of sorbitol and fructose (Fig. 2g). Interestingly, also in the human failing heart, sorbitol release was detected by arteriovenous metabolomics^3^. AKR1B1, a key enzyme of polyol pathway, is also involved in detoxification of reactive aldehydes generated by lipid peroxidation, and its deficiency has been shown to worsen hypertrophy and cardiac dysfunction^27,28^. In contrast, increased flux via the polyol pathway reduces levels of NADPH and increases those of NADH, thereby inhibiting glycolysis, competing with glutathione reductase, increasing ROS, and inhibiting sarcoplasmic/endoplasmic reticulum calcium ATPase^29,30^. Collectively, our data reveal extensive structural and metabolic remodeling in hypertensive HFpEF. Key components of the electron transport chain are significantly downregulated. In parallel, major metabolic pathways, most notably fatty acid oxidation, are suppressed. A compensatory upregulation of glucose import is observed, yet this does not translate into enhanced glycolytic flux. Instead, increased activity of the polyol pathway suggests a diversion of glucose metabolism toward alternative pathways (Extended Data Fig. 3).

From an evolutionary perspective, vital processes such as cardiac energy supply must be prioritized and supported by metabolic flexibility. The heart has an exceptionally high energy demand and must continuously produce large quantities of ATP to maintain its contractile function. Ketone bodies, including β-hydroxybutyrate (BOH), are actively taken up by the heart and can serve as efficient fuel, particularly in HFrEF, where utilization of conventional substrates like long-chain FA is impaired^1,3^. In HFrEF, the ketone-specific contribution to ATP production is approximately 2.5-fold higher than in HFpEF^3^. In untreated hypertensive dTGR HFpEF hearts, BOH accumulated (Fig. 3a), while acetoacetate (AcAc) levels were reduced (Fig. 3a) compared to healthy WT heart. The AcAc-to-BOH ratio (Fig. 3b) indicates an about 40% reduced ketone body oxidation. While BOH infusion improves cardiac output, ejection fraction, and myocardial perfusion in patients with HFrEF^31^, the downregulation of mitochondrial ETC and complex I in dTGR mark a limited metabolization of mitochondrial fuels such as ketone bodies. Thus, we asked whether blood levels of BOH might predict HFpEF disease severity and outcome. Of note, among 23,255 non-fasting participants in the UK Biobank (Table 1), who fulfilled clinical HFpEF criteria following a validated multi-stage algorithm conceptually aligned with ESC recommendations^32^, higher circulating BOH concentrations, quantified by NMR metabolomics, were independently associated with an increased risk of all-cause mortality (HR 1.17, 95% CI 1.14–1.21, p<0.001), consistent with previous observations^33^. When stratified by BOH concentration percentiles (Table 2), mortality risk increased progressively, with significantly lower survival in the highest (>75th percentile) compared with the lowest (<25th percentile) BOH group (log-rank test: χ²=76.86, p<0.001 for non-hypertensive and χ²=84.1, p<0.001 for hypertensive participants; Fig. 3c). In a Cox proportional hazard model for male participants, this association was more pronounced in hypertensive individuals (n=4,287; HR 1.32, 95% CI 1.24–1.40, p<0.001; Fig. 3d) than in non-hypertensive participants (n=9,746; HR 1.13, 95% CI 1.08–1.18, p<0.001; Fig. 3e).

**Fig. 3.**
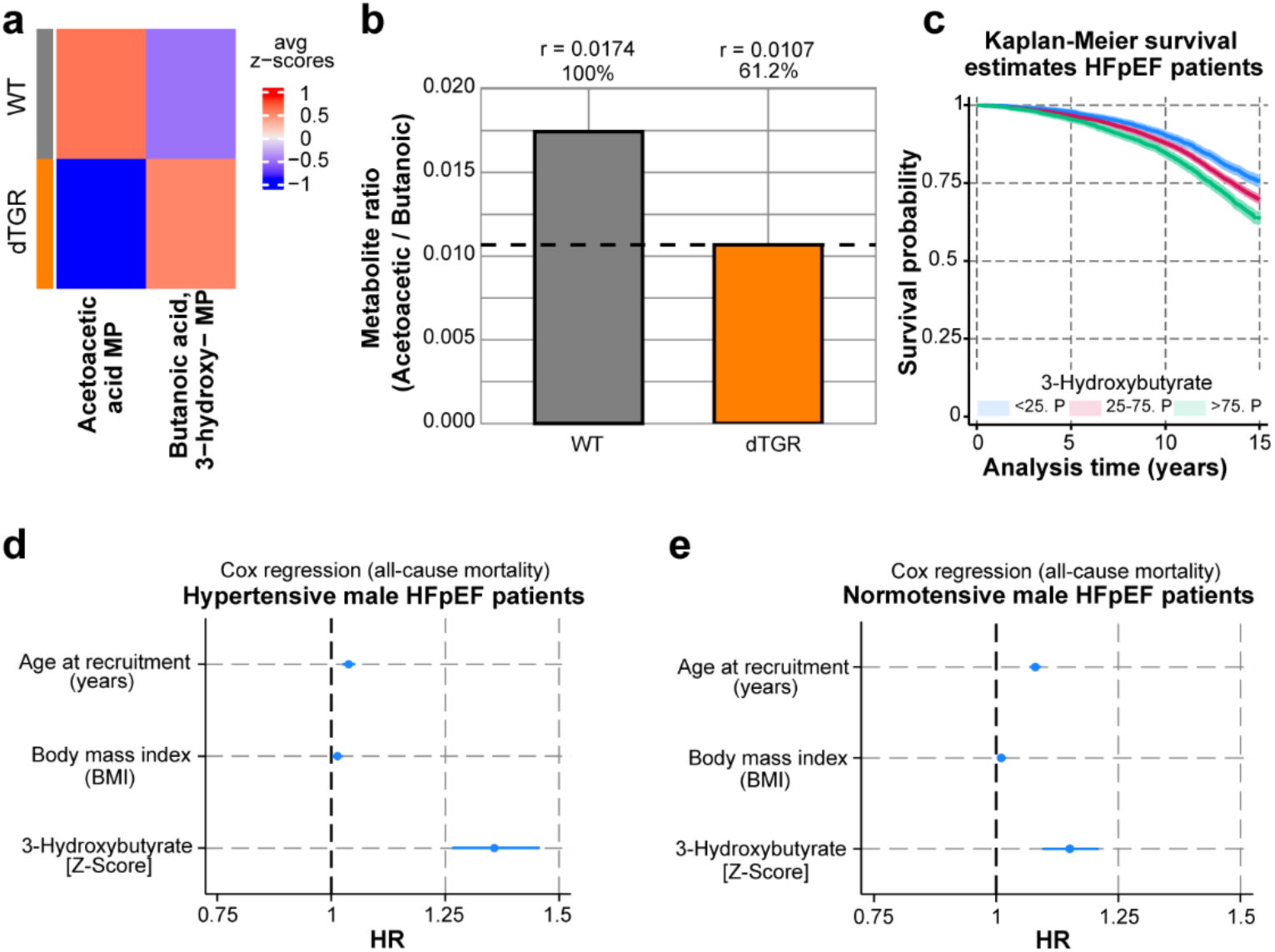
Associations of circulating 3-hydroxybutyrate (BOH) with all-cause mortality in HFpEF patients. **(a)** Heatmap with z-score transformed acetoacetic acid and 3-hydroxy-butanoic acid levels measured by gas chromatography-mass spectrometry. **(b)** Metabolites’ ratio (r) of acetoacetic acid to 3-hydroxy-butanoic acid for WT and dTGR groups accordingly. The percentage for the dTGR group is calculated relative to the WT group, with a dashed line showing the difference between the two groups. **(c)** Kaplan-Meier survival estimates across tertiles of circulating BOH concentrations (<25th, 25th–75th, and >75th percentile) among male HFpEF patients. Patients in the highest 3-hydroxybutyrate tertile (>75th percentile) exhibited the lowest survival probability during follow-up. **(d, e)** Hazard ratios (HRs) with 95% confidence intervals (CIs) derived from Cox regression models (Breslow method for ties), stratified by hypertension status in male HFpEF patients. The proportional hazards assumption was verified and satisfied. **(d)** In the hypertensive subgroup, similar associations were observed (age: HR 1.04 [1.02–1.05], p<0.001; BMI: HR 1.01 [1.00–1.03], p=0.019). BOH concentrations were logarithmically transformed and standardized (Z-score). **(e)** In the normotensive subgroup, age and BMI were positively associated with all-cause mortality (age: HR 1.08 [1.07–1.09], p<0.001; BMI: HR 1.01 [1.00–1.02], p=0.027)

**Table 1.** Clinical characteristics of patients with HFpEF divided into percentiles of circulating 3-hydroxybutyrate.

|  | <25th Perc<br>(N=4,909) | 25th-75th Perc<br>(N=13,256) | >75th Perc<br>(N=5,090) | Total<br>(N=23,255) | P-Value |
| --- | --- | --- | --- | --- | --- |
| Age at recruitment (years) | 62.94 (5.38) | 63.26 (5.13) | 63.70 (4.75) | 63.29 (5.11) | <0.001 |
| Sex | 2,638 (53.7%) | 8,387 (63.3%) | 3,008 (59.1%) | 14,033 (60.3%) | <0.001 |
| Female before menopause | 132 (2.7%) | 221 (1.7%) | 95 (1.9%) | 448 (1.9%) | <0.001 |
| Waist circumference (cm) | 98.40 (13.39) | 102.48 (13.65) | 101.32 (15.31) | 101.37 (14.07) | <0.001 |
| Standing height (cm) | 169.29 (9.46) | 170.27 (9.44) | 169.21 (9.45) | 169.83 (9.46) | <0.001 |
| Seated height (cm) | 137.65 (7.00) | 138.21 (6.99) | 137.15 (7.35) | 137.86 (7.08) | <0.001 |
| Body mass index (BMI) | 30.37 (5.19) | 31.39 (5.42) | 30.96 (5.87) | 31.08 (5.49) | <0.001 |
| Systolic blood pressure (mmHg) | 143.31 (19.24) | 143.73 (19.12) | 146.25 (19.41) | 144.19 (19.24) | <0.001 |
| Diastolic blood pressure (mmHg) | 82.03 (10.64) | 82.63 (10.69) | 83.75 (10.60) | 82.75 (10.67) | <0.001 |
| Pulse rate (bpm) | 67.74 (11.91) | 69.52 (13.15) | 72.39 (13.81) | 69.77 (13.14) | <0.001 |
| Whole body fat mass (kg) | 29.99 (10.58) | 31.49 (11.00) | 31.07 (11.74) | 31.08 (11.10) | <0.001 |
| Whole body fat-free mass (kg) | 57.15 (11.96) | 59.62 (11.96) | 57.66 (12.18) | 58.67 (12.06) | <0.001 |
| Whole body water mass (kg) | 41.85 (8.76) | 43.66 (8.77) | 42.23 (8.93) | 42.96 (8.84) | <0.001 |
| Pulse pressure (mmHg) | 61.28 (15.22) | 61.10 (15.27) | 62.48 (15.81) | 61.44 (15.39) | <0.001 |
| Appendicular muscle mass (kg) | 24.23 (5.62) | 25.50 (5.78) | 24.64 (5.92) | 25.04 (5.80) | <0.001 |
| Whole body fat mass (%) | 0.34 (0.09) | 0.34 (0.08) | 0.34 (0.09) | 0.34 (0.09) | 0.061 |
| Whole body fat-free mass (%) | 0.66 (0.09) | 0.66 (0.09) | 0.66 (0.09) | 0.66 (0.09) | 0.076 |
| Whole body water mass (%) | 0.48 (0.06) | 0.48 (0.06) | 0.48 (0.06) | 0.48 (0.06) | 0.068 |
| Death (all causes) | 1,074 (21.9%) | 3,570 (26.9%) | 1,633 (32.1%) | 6,277 (27.0%) | <0.001 |
| Postviral fatigue | 19 (0.4%) | 55 (0.4%) | 19 (0.4%) | 93 (0.4%) | 0.911 |
| Sleep apnoea | 67 (1.4%) | 235 (1.8%) | 68 (1.3%) | 370 (1.6%) | 0.038 |
| Chronic ischaemic heart disease | 682 (13.9%) | 2,088 (15.8%) | 666 (13.1%) | 3,436 (14.8%) | <0.001 |
| Nonrheumatic mitral valve disease | 88 (1.8%) | 219 (1.7%) | 81 (1.6%) | 388 (1.7%) | 0.716 |
| Nonrheumatic aortic valve disease | 70 (1.4%) | 188 (1.4%) | 81 (1.6%) | 339 (1.5%) | 0.667 |
| Cardiomyopathy | 48 (1.0%) | 133 (1.0%) | 46 (0.9%) | 227 (1.0%) | 0.828 |
| Varicose veins of lower extremities | 156 (3.2%) | 316 (2.4%) | 106 (2.1%) | 578 (2.5%) | 0.001 |
| Hypotension | 54 (1.1%) | 127 (1.0%) | 58 (1.1%) | 239 (1.0%) | 0.470 |
| Angina pectoris/ coronary artery disease | 506 (10.3%) | 1,568 (11.8%) | 498 (9.8%) | 2,572 (11.1%) | <0.001 |
| Endocrine, nutritional and metabolic diseases | 1,023 (20.8%) | 3,221 (24.3%) | 1,254 (24.6%) | 5,498 (23.6%) | <0.001 |
| Mental and behavioural disorders | 180 (3.7%) | 513 (3.9%) | 227 (4.5%) | 920 (4.0%) | 0.094 |
| Diseases of the nervous system | 445 (9.1%) | 1,319 (10.0%) | 463 (9.1%) | 2,227 (9.6%) | 0.083 |
| Diseases of the eye and adnexa | 395 (8.0%) | 1,084 (8.2%) | 433 (8.5%) | 1,912 (8.2%) | 0.676 |
| Diseases of the respiratory system | 658 (13.4%) | 1,997 (15.1%) | 820 (16.1%) | 3,475 (14.9%) | <0.001 |
| Diseases of the digestive system | 1,575 (32.1%) | 4,531 (34.2%) | 1,697 (33.3%) | 7,803 (33.6%) | 0.027 |
| Diseases of the skin and subcutaneous tissue | 439 (8.9%) | 1,215 (9.2%) | 475 (9.3%) | 2,129 (9.2%) | 0.795 |
| Diseases of the musculoskeletal system and connective tissue | 1,258 (25.6%) | 3,333 (25.1%) | 1,300 (25.5%) | 5,891 (25.3%) | 0.744 |
| Diseases of the genitourinary system | 1,027 (20.9%) | 2,659 (20.1%) | 988 (19.4%) | 4,674 (20.1%) | 0.167 |
| Pregnancy, childbirth and the puerperium | 23 (0.5%) | 19 (0.1%) | 7 (0.1%) | 49 (0.2%) | <0.001 |
| Congenital malformations, deformations and chromosomal abnormalities | 72 (1.5%) | 193 (1.5%) | 72 (1.4%) | 337 (1.4%) | 0.972 |
| Aortic aneurysm | 24 (0.5%) | 83 (0.6%) | 31 (0.6%) | 138 (0.6%) | 0.557 |
| Phlebitis and thrombophlebitis | 139 (2.8%) | 350 (2.6%) | 146 (2.9%) | 635 (2.7%) | 0.619 |
| Type 2 diabetes | 312 (6.4%) | 1,553 (11.7%) | 777 (15.3%) | 2,642 (11.4%) | <0.001 |
| Myocardial infarction | 349 (7.1%) | 1,073 (8.1%) | 323 (6.3%) | 1,745 (7.5%) | <0.001 |
| Chronic kidney disease | 271 (5.5%) | 869 (6.6%) | 306 (6.0%) | 1,446 (6.2%) | 0.029 |
| Hypertensive renal disease | 37 (0.8%) | 103 (0.8%) | 56 (1.1%) | 196 (0.8%) | 0.075 |
| Endocarditis | 50 (1.0%) | 122 (0.9%) | 47 (0.9%) | 219 (0.9%) | 0.821 |
| Essential (primary) hypertension | 1,320 (26.9%) | 4,047 (30.5%) | 1,501 (29.5%) | 6,868 (29.5%) | <0.001 |
| Atrial fibrillation and flutter | 961 (19.6%) | 2,355 (17.8%) | 792 (15.6%) | 4,108 (17.7%) | <0.001 |
| Cerebrovascular event/ stroke | 187 (3.8%) | 519 (3.9%) | 199 (3.9%) | 905 (3.9%) | 0.945 |
| Embolism and thrombosis | 235 (4.8%) | 642 (4.8%) | 248 (4.9%) | 1,125 (4.8%) | 0.980 |
| Pulmonary embolism | 78 (1.6%) | 170 (1.3%) | 57 (1.1%) | 305 (1.3%) | 0.108 |
| Pulmonary artery hypertension | 19 (0.4%) | 51 (0.4%) | 17 (0.3%) | 87 (0.4%) | 0.868 |
| Neoplasms | 918 (18.7%) | 2,630 (19.8%) | 1,059 (20.8%) | 4,607 (19.8%) | 0.030 |
| Congenital malformations of the circulatory system | 34 (0.7%) | 63 (0.5%) | 34 (0.7%) | 131 (0.6%) | 0.117 |
| Seizure/Epilepsy | 93 (1.9%) | 184 (1.4%) | 80 (1.6%) | 357 (1.5%) | 0.047 |
| Mineralocorticoid receptor antagonist (MRA) | 34 (0.7%) | 148 (1.1%) | 56 (1.1%) | 238 (1.0%) | 0.035 |
| Levothyroxine | 230 (4.7%) | 509 (3.8%) | 231 (4.5%) | 970 (4.2%) | 0.014 |
| Metformin | 111 (2.3%) | 695 (5.2%) | 411 (8.1%) | 1,217 (5.2%) | <0.001 |
| Warfarin | 310 (6.3%) | 758 (5.7%) | 237 (4.7%) | 1,305 (5.6%) | 0.001 |
| Sulfonylurea | 66 (1.3%) | 370 (2.8%) | 225 (4.4%) | 661 (2.8%) | <0.001 |
| Iron therapy | 232 (4.7%) | 544 (4.1%) | 228 (4.5%) | 1,004 (4.3%) | 0.152 |
| Beta blocker | 1,976 (40.3%) | 5,372 (40.5%) | 1,597 (31.4%) | 8,945 (38.5%) | <0.001 |
| ACE inhibitor | 1,711 (34.9%) | 5,404 (40.8%) | 2,109 (41.4%) | 9,224 (39.7%) | <0.001 |
| Angiotensin receptor blocker (ARB) | 694 (14.1%) | 2,215 (16.7%) | 769 (15.1%) | 3,678 (15.8%) | <0.001 |
| Loop diuretic | 481 (9.8%) | 1,521 (11.5%) | 605 (11.9%) | 2,607 (11.2%) | 0.001 |
| Calcium channel blocker | 1,294 (26.4%) | 4,127 (31.1%) | 1,599 (31.4%) | 7,020 (30.2%) | <0.001 |
| Aspirin | 1,920 (39.1%) | 5,875 (44.3%) | 2,108 (41.4%) | 9,903 (42.6%) | <0.001 |
| Thiazide diuretic | 1,176 (24.0%) | 3,646 (27.5%) | 1,405 (27.6%) | 6,227 (26.8%) | <0.001 |
| Cholesterol lowering medication | 2,027 (41.3%) | 6,410 (48.4%) | 2,254 (44.3%) | 10,691 (46.0%) | <0.001 |
| Insulin | 87 (1.8%) | 505 (3.8%) | 286 (5.6%) | 878 (3.8%) | <0.001 |

**Table 2.**
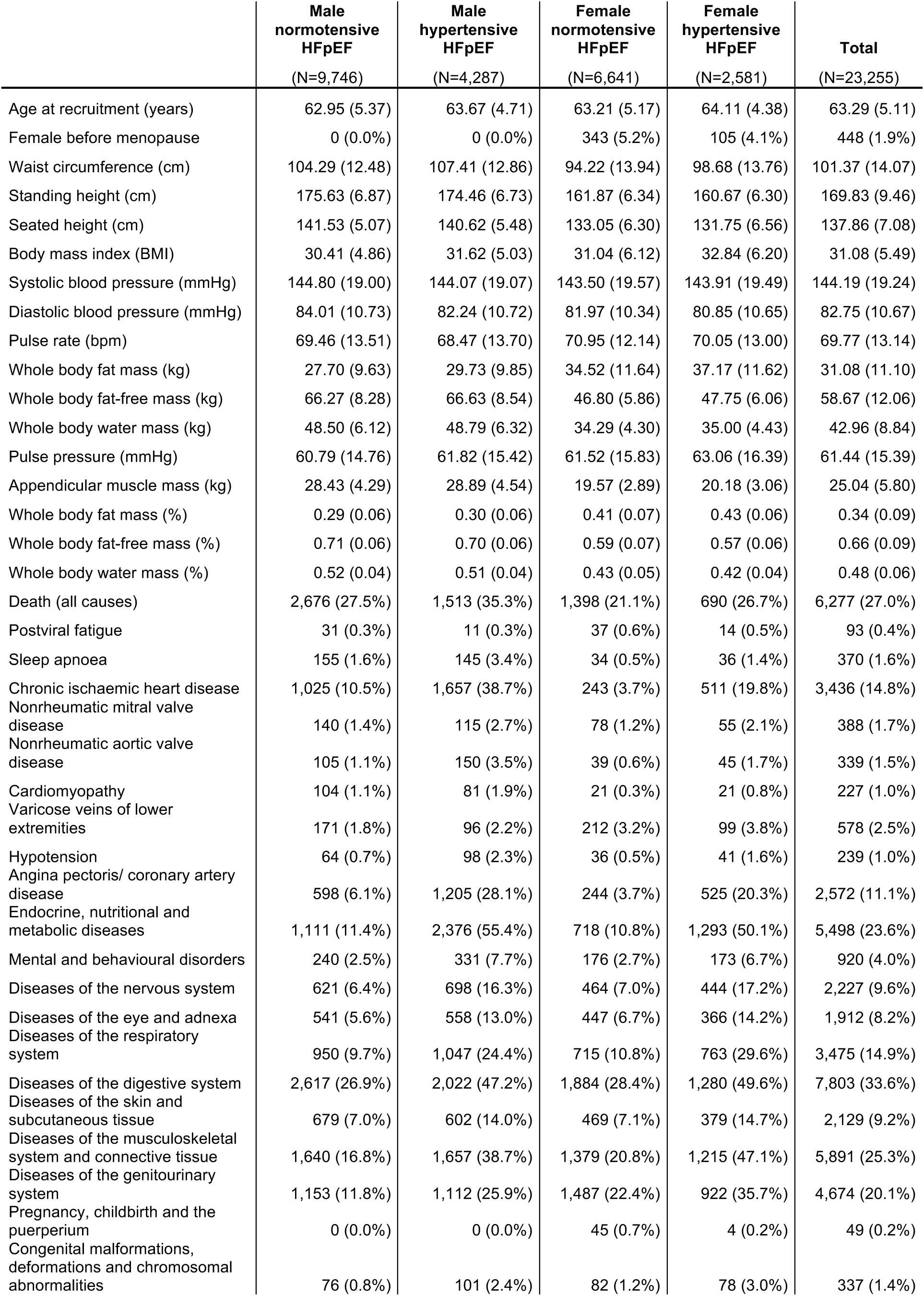
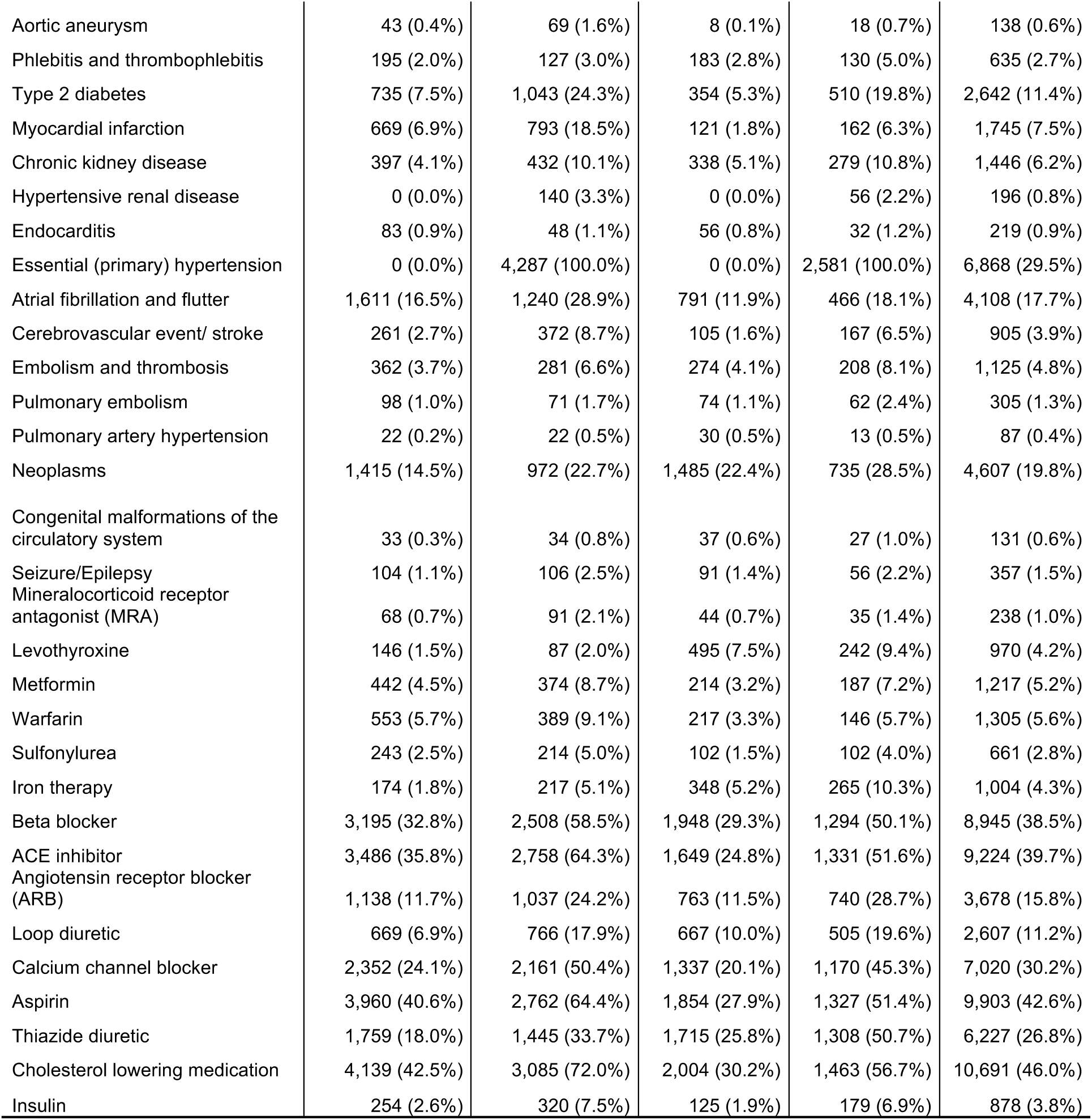
Demographic characteristics, comorbidities, and medication profiles across hypertensive and non-hypertensive HFpEF subgroups, separated by sex.

We do not consider BOH to be a classic risk factor for HFpEF - but argue that increased BOH levels indicate, that not substrate availability, but limited mitochondrial metabolism due to downregulation of electron transport chain proteins cause its increase. We hypothesize that BOH accumulates in the circulation, because BOH is not effectively processed into AcAc due to reduced NAD^+^ levels. This is consistent with early biochemical studies by Hans Krebs demonstrating that NAD^+^ to NADH ratio directly reflects BOH to AcAc imbalance^34^. In general, reduced BOH metabolism into acetyl-CoA suggests that BOH cannot be fully utilized for ATP production, meaning it cannot compensate for the energy deficit in HFpEF hearts. This may explain the increased risk in HFpEF patients.

### Butyrate serves as alternative fuel and improves diastolic dysfunction

Given the biochemical relationship between butyrate and BOH, which both enter the mitochondrial β-oxidation pathway of fatty acids at a late stage, and the fact that butyrate is an established primary energy source for colonocytes, we hypothesized that butyrate may also act as an additional ancillary cardiac fuel in HFpEF. Butyrate is typically produced in the colon by SCFA-producing bacteria. After absorption by the host, butyrate is either immediately oxidized by colonocytes to support their energy needs or transferred to the bloodstream via interstitial and lymph fluid. Interestingly, dTGR rats had similar colonic fecal butyrate concentrations compared to WT controls indicating a comparable microbial SCFA production (Fig. 4a). However, compared to WT, butyrate levels in the interstitial fluid of the colon and circulation were significantly lower in hypertensive dTGR HFpEF rats (Fig. 4b, c), consistent with observations in the circulation of HFpEF patients^9^. The reduced butyrate levels in the host suggest that butyrate is consumed locally by epithelial cells and other metabolically active cells, such as those in the heart and kidneys. To directly test this hypothesis, we performed metabolic tracing and functional analyses as well as chronic butyrate treatment in our HFpEF model. In Seahorse experiments of healthy rat cardiomyocytes, butyrate significantly increased basal oxidative respiration exceeding that of BOH and amino acids and reaching levels comparable to those of long-chain fatty acids (FA) (Fig. 4d). Further we applied ¹³C-butyrate tracing in isolated perfused hearts from WT and dTGR HFpEF animals to follow its myocardial utilization *in situ*. We observed ^13^C-isotope incorporation in butyryl-CoA and downstream metabolites confirming active substrate usage in heart. When comparing isotope incorporation in butyryl-CoA (ButCoA), ^13^C-isotope incorporation was increased in dTGR compared to WT hearts (Fig. 4e).

**Fig. 4.**
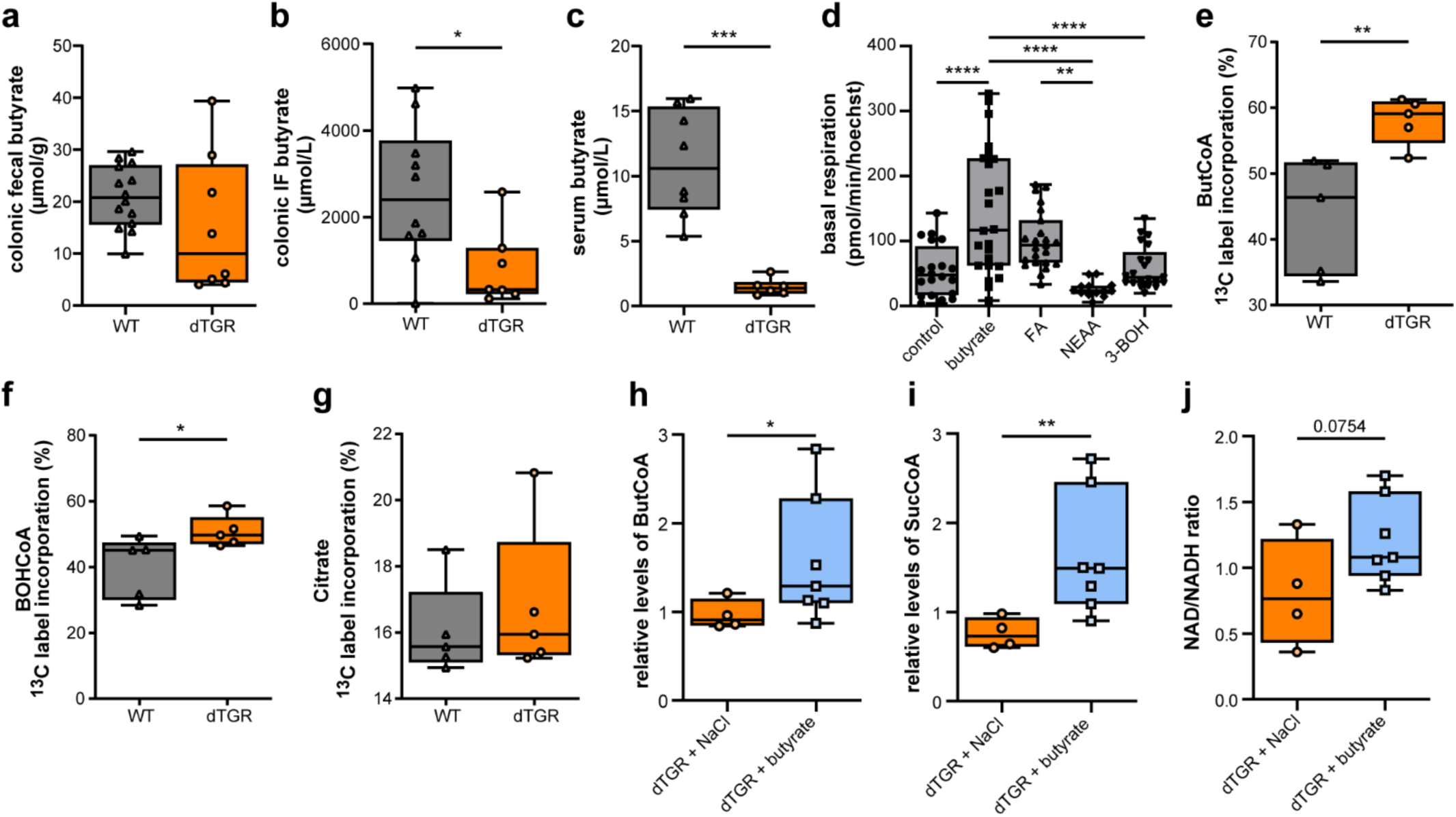
Impaired systemic butyrate supply but enhanced cardiac utilization in HFpEF, with metabolic rescue by chronic butyrate supplementation. **(a)** Colon feces butyrate levels in µmol/g wet weight measured by gas chromatography-mass spectrometry (GC-MS) (n=14 WT, n=8 dTGR). **(b)** Colonic interstitial fluid (IF) butyrate levels in µmol/L measured by GC-MS (n=10 WT, n=7 dTGR). **(c)** Serum butyrate levels in µmol/L measured by GC-MS (n=8 WT, n=6 dTGR). **(d)** Basal oxidative respiration measured as pmol/min/hoechst intensity in isolated healthy rat cardiomyocytes treated with glucose and glutamine only (control n=20) or supplemented with butyrate (n=24), fatty-acids (FA n=23), non-essential amino acids (NEAA n=13) or beta-hydroxybutyrate (BOH n=21) in addition to glucose and glutamine. **(e-g)** Natural abundance corrected ^13^C isotope incorporation (indicated as percentage of labeled in total) in butyryl-CoA **(e)**, BOH-CoA **(f)**, and citrate **(g)** in ex vivo perfused WT and dTGR hearts (n=5 WT, n=5 dTGR). **(h-j)** Relative metabolite levels of butyryl-CoA **(h)**, succinyl-CoA **(i)**, and the NAD/NADH ratio **(j)** in dTGR treated with NaCl (n=4) or butyrate (n=7) hearts. Data are presented as boxplots (IQR) with whiskers min to max. **(b)** Technical replicates pooled from two independent experiments. **(a-c)** *P<0.05, ***P<0.001 unpaired two-tailed Welch’s t-test; **(d)** **P<0.01, ****P<0.0001 one-way ANOVA with Tukey’s multiple comparison. **(e-j)** *P<0.05, **P<0.01 unpaired one-tailed Welch’s t-test.

Next, we investigated whether the chronic butyrate treatment would affect the metabolism in hypertensive HFpEF hearts by using it as an energy substrate. For three weeks, dTGR rats received either 200 mM sodium-chloride (dTGR+NaCl) or 200 mM sodium-butyrate (dTGR+butyrate) via drinking water. Non-transgenic NaCl-treated WT served as controls (WT+NaCl). Equimolar sodium was used to control the sodium load, since in humans and rodents, high sodium intake is a risk factor for hypertension and CVD. Metabolic profiling of dTGR upon chronic butyrate treatment resulted in significantly increased cardiac butyrylCoA (Fig. 4h), succinylCoA (Fig. 4i) levels with a trend toward an increased NAD/NADH ratio (Fig. 4j). The competence of healthy and dTGR HFpEF heart muscle to directly metabolize butyrate and the observed rewinding of the metabolic phenotype upon chronic butyrate supplementation suggest a direct participation of butyrate in cardiac metabolism. Therefore, we hypothesized that chronic butyrate treatment not only resulted in an improved cardiac energy metabolism, but also functionally improved diastolic dysfunction. NaCl-treated dTGR showed an aggravated HFpEF progression with high mortality (Fig. 5a). NaCl-treated dTGR exhibited high mortality (Fig. 5a) and progressive disease. In contrast, Na-butyrate significantly improved survival and stabilized body weight (Fig. 5a; Extended Data Fig. 4a). However, both dTGR groups were severely hypertensive, as determined by tail-cuff and continuous radiotelemetry recordings (Fig. 5b-c). Na-butyrate did not attenuate cardiac hypertrophy, as indicated by posterior wall thickness (Fig. 5d), heart weight-to-tibia length ratio (Extended Data Fig. 4b), or cardiomyocyte cross-sectional area (WGA staining, Extended Data Fig. 4c) most likely a consequence of the severe hypertension (>200 mmHg).

**Fig. 5.**
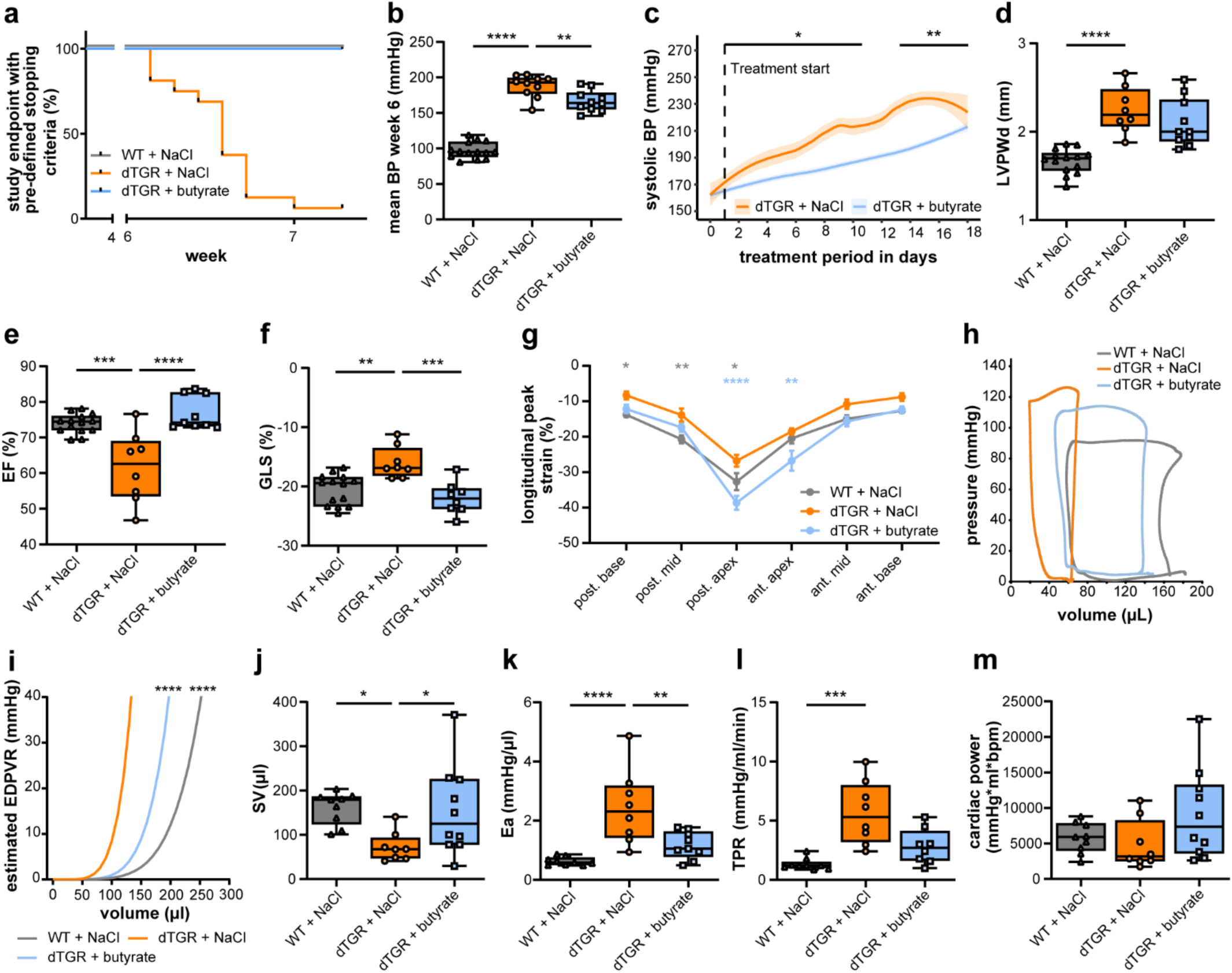
SCFA Na-butyrate mitigates myocardial pathology in hypertensive HFpEF despite persisting hypertrophy. **(a)** Kaplan Meier curve of survival represents percentage of animals reaching study endpoint. (n=10-16) **(b)** Mean blood pressure (BP) at week 6 measured by tail cuff method. (n=10-14) **(c)** Systolic BP was measured continuously by radiotelemetry in dTGR treated with NaCl (n=6) or butyrate (n=5). To assess butyrate’s effect on SBP, we used linear mixed-effects models on 5-minute SBP data. Adding medication (dTGR+butyrate vs dTGR+NaCl) as a fixed effect significantly improved model fit (χ²(1)=15.06, p=0.0001), indicating a treatment effect. Daily group differences were identified via unpaired t-tests on mean SBP per animal. **(d)** Cardiac hypertrophy assessed in echocardiography through LV posterior wall thickness in diastole (LVPWd, n=8-14). **(e)** Ejection fraction (EF) measured by echocardiography (n=8-14). **(f)** Mean global longitudinal strain (GLS) was assessed in parasternal long axis (PLAX) B-mode echocardiography (n=8-14). **(g)** Segmental longitudinal peak strain in PLAX (n=8-14). **(h)** Representative LV ventricular pressure-volume loops measured with the Millar Tip catheter system are displayed for one animal per group. **(i)** Estimated EDPVR (n=7-9), **(j)** stroke volume (SV) **(k)**, arterial elastance (Ea) (n=8-9), and **(l)** total peripheral resistance (TPR, n=8-9). **(m)** Cardiac power (mmHg*ml*bpm) was similar in WT and dTGR (n=8-10). Outliers were removed upon statistical testing. Data are presented as boxplots (IQR) with whiskers min to max. **(b, d-f and j-m)**. *P<0.05, **P<0.01, ***P<0.001, ****P<0.0001 **(b, d, e, f, j, k)** via ordinary one-way ANOVA with Dunnett’s multiple comparison, **(l, m)** Kruskal-Wallis test with Dunn’s multiple comparison, **(g, h)** two-way ANOVA with Dunnett‘s multiple comparison.

Importantly, Na-butyrate significantly ameliorated diastolic dysfunction. Ejection fraction (EF) was preserved in Na-butyrate-treated dTGR and reached levels comparable to healthy controls (Fig. 5e). Global longitudinal strain (GLS), was significantly improved in Na-butyrate-treated dTGR (Fig. 5f) with regional recovery most pronounced in the posterior apex (Fig. 5g), the same region previously identified as showing the most prominent structural and proteomic changes. Invasive hemodynamic analysis demonstrated partial normalization of pressure-volume loop geometry (Fig. 5h) and improved diastolic compliance, reflected by a rightward shift of the end-diastolic pressure-volume relationship (EDPVR, Fig. 5i) after Na-butyrate treatment. Stroke volume was significantly increased (Fig. 5j), and both arterial elastance (Ea, Fig. 5k) and total peripheral resistance (TPR, Fig. 5l) were reduced, indicating improved ventricular-vascular coupling and overall hemodynamic performance. Cardiac power was preserved, with a modest increase observed in Na-butyrate-treated dTGR (Fig. 5m). Along this line, the expression of key molecular markers of remodeling, including atrial natriuretic peptide (Anp), brain natriuretic peptide (Bnp), TIMP metallopeptidase inhibitor 1 (Timp1), and osteopontin (Opn) were significantly decreased (Extended Data Fig. 4d-g), indicating transcriptional attenuation of pathological remodeling. Histological analysis confirmed these molecular findings. Na-butyrate reduced perivascular fibrosis (Sirius Red staining, Extended Data Fig. 4h) and CD68-positive macrophage abundance (Extended Data Fig. 4i), consistent with reduced fibrotic and inflammatory burden. Titin isoforms in the myocardium are known to reflect the myofilament stiffness. Our analysis revealed no significant difference in N2BA/N2B titin isoform ratio between NaCl-treated and Na-butyrate groups (data not shown), suggesting that other mechanisms are responsible for cardiac improvement.

Together, these findings demonstrate that chronic Na-butyrate supplementation ameliorates metabolic, functional and structural features of hypertensive HFpEF, despite persistent hypertension and cardiac hypertrophy.

## Discussion

Even in the advanced stages of the disease, the failing heart of our hypertensive HFpEF model maintains a cardiac output similar to that of a healthy non-failing heart, despite an obvious imbalance between energy demand and supply. Our ^13^C-butyrate tracing experiments suggest that butyrate enters the TCA cycle as a metabolically accessible substrate, counterregulating failing oxidative metabolism and supporting diastolic function. These findings reveal a new non-hemodynamic, substrate-based therapeutic axis.

Current treatment strategies for HFpEF are limited. Unlike HFrEF, where therapies targeting neurohormonal and hemodynamic pathways consistently improve outcomes, most interventions that are effective for systolic heart failure have failed for HFpEF. This discrepancy likely reflects the systemic, multiorgan nature of HFpEF. The first therapy shown to consistently benefit HFpEF patients was the sodium-glucose cotransporter 2 (SGLT2) inhibitor. Their pleiotropic action has been linked to the “thrifty substrate hypothesis,” which proposes that circulating BOH serves as an efficient fuel under metabolic stress. Indeed, SGLT2 inhibitors increase BOH levels, and acute BOH infusion improves cardiac output, ejection fraction, and myocardial perfusion in patients with HFrEF without compromising oxygen economy^33^. However, our experimental data in dTGR and the observational findings from the UK Biobank (Fig. 2f, 3a–e) point to a metabolic bottleneck in BOH handling in HFpEF. This is reflected by BOH accumulation and a reduced AcAc-to-BOH ratio (Fig. 3a), indicating redox imbalance and impaired oxidative capacity. Our findings of butyrate’s efficacy point to an approach complementary to SGLT2 inhibition by fueling the cardiac metabolic substrate pool.

From a biochemical perspective, BOH and butyrate converge on the mitochondrial acetyl-CoA pool through mechanistically distinct entry points. BOH must first be oxidized to AcAc by β-hydroxybutyrate dehydrogenase, a fully reversible reaction governed by the mitochondrial NAD⁺/NADH ratio, and is then activated to acetoacetyl-CoA via succinyl-CoA:3-oxoacid CoA transferase, which exchanges CoA between succinyl-CoA and acetoacetate without direct ATP consumption and operates close to equilibrium. In contrast, butyrate is activated by mitochondrial AMP-forming acyl-CoA synthetases to butyryl-CoA in an ATP-dependent reaction that yields AMP and pyrophosphate; rapid pyrophosphate hydrolysis renders this step essentially irreversible and strongly biased towards forward flux. Subsequent β-oxidation of butyryl-CoA produces acetyl-CoA independently of the redox constraints that limit ketone oxidation in the setting of impaired electron transport chain function. These fundamentally different enzymatic properties provide a mechanistic rationale for why butyrate, despite its lower in vivo concentrations, can be efficiently funneled into mitochondrial oxidation and support cardiac energetics in HFpEF, whereas BOH accumulates when redox balance and TCA cycle flux are constrained.

Limitations of our study include its reliance on a single transgenic model that predominantly reflects the hypertensive phenome of HFpEF and therefore does not encompass the clinical heterogeneity of the human syndrome. Yet this constraint also represents a conceptual strength: by isolating the hypertensive component of HFpEF pathophysiology, the model allows us to dissect the mitochondrial derangements that arise specifically from chronic pressure overload.

In this context, our multi-omic data and the UK Biobank analysis converge to show that elevated circulating BOH indicates reduced ketone utilization. Rather, it points to failed ketone oxidation and mitochondrial bottlenecks, which are associated with increased mortality. These observations reinforce the notion that mitochondrial dysfunction and insufficient BOH usage are integral features of the hypertensive HFpEF phenotype across species. Long-term safety, dose optimization and the impact of comorbidities common in HFpEF (e.g., obesity, diabetes, chronic kidney disease) will require dedicated evaluation, and we cannot exclude that butyrate’s benefits may extend beyond its energetic role to include epigenetic or immunomodulatory actions.

In summary, our findings position butyrate as a simple, readily utilizable “rescue fuel” capable of bypassing redox-sensitive metabolic bottlenecks in the pressure-overloaded heart. Other energy-depleted states, such as myocardial ischemia, sepsis, or inherited mitochondrial disorders, may similarly profit from strategies that raise circulating butyrate levels. Moreover, the profound metabolic impairment of donor organs during cold preservation suggests that butyrate supplementation could improve organ viability and post-transplant function. These hypotheses warrant rigorous testing in randomized clinical trials in patients with HFpEF and in controlled studies of organ procurement and preservation.

## Methods

### Animal ethics, breeding and housing

The animal study was registered and approved by the local ethics board (G0036/20) and conducted according to the German/European law for animal protection. A detailed ARRIVE 2.0 compliance checklist for this study is provided in Extended Table 1. All animals were housed in the facility of the Max-Delbrück-Center for Molecular Medicine (MDC). Female Sprague Dawley rats homozygous transgenic for renin (SD-Tg(hREN)L10J) were mated with male Sprague Dawley rats homozygous transgenic for angiotensinogen (SD-Tg(hAGT)L1623)^18^. Male double transgenic rats (dTGR), expressing both human renin and angiotensinogen, entered the study at the age of four weeks. Age-matched non-transgenic wild-type Sprague Dawley control rats (WT) were included into the study. All dTGR and WT breeding was conducted at the animal facility of MDC. Animals were housed in small groups (2-5/cage) under controlled standard conditions with a 12/12 h light/dark cycle, temperature of 22 ± 2°C and humidity of 55 ± 10%. During telemetry admission animals were single-housed. Water and standard chow (Sniff V1324-300) were provided ad libitum throughout the experiment.

### Animal protocols

Animals were randomized to groups with regards to litter and body weight. One study group consisted of untreated dTGR and untreated WT. A second study group received either 200 mM sodium-chloride (NaCl treated WT and NaCl treated dTGR) or 200 mM sodium-butyrate (Na-butyrate treated dTGR) via drinking water. After three weeks of treatment, animals were sacrificed at the age of 7 weeks. In accordance with the 3Rs principle (Replace, Reduce, Refine), all animal experiments were designed to use the minimal number of animals required, lower the potential distress to a minimal level, and incorporate alternatives when feasible.

Animals were used for (I) functional phenotyping including weekly tail-cuff blood pressure assessment, echocardiography at the age of 6 weeks and sacrifice at 7 weeks of age with myocardial analysis post mortem (cohort I: WT n=13; dTGR n=13, cohort II NaCl treated WT n=14; NaCl treated dTGR n=16; Na-butyrate treated dTGR n=10); (II) hemodynamics with Millar tip catheter measurements at the age of 6 weeks with subsequent sacrifice (cohort II: WT n=10; dTGR n=8; cohort II NaCl treated WT n=10; NaCl treated dTGR n=9; Na-butyrate treated dTGR n=10); and (III) continuous blood pressure cohort with telemetry transmitter implantation at 3 weeks of age and blood pressure assessment until sacrifice at the age of 7 weeks (cohort III: NaCl treated dTGR n=6; Na-butyrate treated dTGR n=5). Animals were sacrificed under isoflurane anesthesia (3-4%), serum and plasma were collected and heart tissue was harvested, snap frozen in liquid nitrogen for RNA analysis and stored at −70°C until further used or fixed in formalin for histology. Rat hearts were dissected in longitudinal or horizontal orientation. The LV was dissected according to the six regions with LV free wall into posterior base, mid, apex and LV septal wall into anterior base, mid, apex according to echocardiography analysis.

### Echocardiography

Rats (cohort I) were weighted, initially anesthetized with 3-4% isoflurane and positioned on a 37°C heat pad where body temperature was controlled throughout the measurement. Anesthesia was maintained through a breathing mask with minimum (1-2.5%) isoflurane. The heart rate and echocardiogram (ECG) were monitored. Hair was removed from the chest and ultrasound gel was applied. Echocardiography scans were recorded in parasternal long and short axis view using the high-resolution ultrasound system Vevo 3100 (Fujifilm, VisualSonics Inc.) with a 30 MHz ultrasound transducer (MX250). All echocardiography parameters were calculated according to the standard protocol (Vevo 770® Standard Measurements and Calculations, Rev 1.2; Visual Sonics) using the Vevo LAB Software. B-Mode analysis was used for systolic and diastolic functional parameter of the LV. M-Mode analyses were used to gain conventional two-dimensional parameters on cardiac function.

Parasternal long axis B-Mode loops were filtered according to their image acquisition quality and used to assess the shortening of the myocardium as global longitudinal strain (GLS) and regional longitudinal peak strain as described before^35^. The endocardium was manually traced during the end-diastolic phase, while the epicardium was automatically captured by the software and only corrected when necessary. The strain was calculated from three independent measurements of three consecutive cardiac cycles. The regional strain was calculated for six segments each defined by eight speckles. The cardiac regions are defined according to standardized myocardial segmentation^36^.

### Hemodynamic measurement

Carprofen (5 mg/kg BW; s.c.) was applied 30 min prior to the initial anesthesia with isoflurane (3-4%). Narcosis was maintained with 1-2% isoflurane. Animals (cohort II) were intubated and kept on a heating plate at 37°C throughout the experiment. Hair was removed, skin disinfected and the Millar tip catheter inserted through the right carotid artery into the LV. To validate the correct placement of the catheter, echocardiography and pressure-volume-values were used. The animals were sacrificed under final narcosis and subsequent organs and blood harvested. Hemodynamic parameters were determined using LabChart software (v8.1.13; AD Instruments). The predicted end-diastolic pressure-volume relationship (EDPVR) was determined according to the method described by Klotz et al^37^. Arterial elastance (Ea) was calculated as Ea = LVESP / (Vmax - Vmin). Total peripheral resistance (TPR) was derived using the formula TPR = MAP / CO. Cardiac power^38,39^ was calculated as the product of stroke volume (SV), mean arterial pressure (MAP), and heart rate (HR).

### Blood pressure assessment

Non-invasive tail cuff blood pressure measurement was carried out at single timepoints at five, six and seven weeks of age (cohort I). The animal was placed in the rodent holder (CODA system), the tail cuff was attached to the tail and blood pressure measured after 5-10 min adaptation. In cohort III, invasive BP measurement was conducted using a telemetric system (DSI). A telemetric transmitter (PA-C10; DSI) was implanted, sewed to the peritoneum and the sensor located infrarenal in the abdominal aorta at the age of three weeks. The abdomen closed with surgical clips. Animals received metamizole one day pre-surgery and three days post-surgery (400 mg/kg/24h) for analgesia. Carprofen (5 mg/kg BW; s.c.) was applied 30 min prior to the initial anesthesia with isoflurane (3-4%). Each cage was placed on its respective transmitter detection plate; therefore single-housing was required. Animals from the telemetry cohort were sacrificed at the age of seven weeks with isoflurane (3-4%) anesthesia, followed by blood and organ harvest.

### Blood analysis

Serum was collected after centrifugation of blood (2500xg, 10 min). For plasma, blood was anticoagulated using Na-EDTA (Sigma) and centrifuged at 1500xg, 15 min, 4°C Both were immediately frozen and stored at −70°C. Plasma levels of brain natriuretic peptide (BNP, abcam ab108816) were detected using enzyme-linked immunosorbent assay (ELISA), according to the manufacturer protocol. Butyrate serum levels were detected using gas chromatography-mass spectrometry as described earlier.^40^

### Interstitial fluid and colon content isolation

The interstitial fluid was isolated from rat colon as described before by Avery et al.^40^ In short, the intestine was dissected into respective segments. Content was removed from colon, snap frozen and stored for further analyzed using gas chromatography-mass spectrometry. Tissue was weight and incubated in 0.9% NaCl at a ratio 1:10 for 2 h at 4°C on a beaker. Resulting interstitial fluid was filtered (0.2 µm) and stored at −70°C until further analyzed using gas chromatography-mass spectrometry.

### Cardiomyocyte isolation, Seahorse assay and ^13^C4 Na-butyrate isotope labeling

Cardiomyocytes from WT (n=7) and dTGR (n=6) were isolated using the Langendorff-free isolation method^41^. In short, the heart was quickly removed after decapitation under isoflurane narcosis and EDTA containing perfusion buffer (pH 7.4) was injected with a cannula through the LV apex. Excess EDTA was removed using perfusion buffer (pH 7.4), following a 30 min digestion with digestion solution containing collagenase II, IV (Worthington LS004176, LS004188) and protease (Sigma P5147). Subsequently, the heart was cut into pieces and pulled apart with forceps. Digestion reaction was terminated using stop buffer, perfusion buffer supplemented with 5% FBS. After 30 min of settling and calcium reintroduction, isolated primary cardiomyocytes were seeded into a 96-well seahorse cell culture plate with a density of 1 x 10^3^/well (4-6 technical replicates per animal). After 45 min settling and 60 min culture in M199 medium (Pan-Biotech P04-07050) containing 2% (v/v) bovine serum albumin (Sigma, A7906), 1% (v/v) Insulin-Transferrin-Selenium (Sigma, I3146, 100x), 1% (v/v) CD lipids (Gibco 11905031, 100%), 1% (v/v) Penicillin/Streptomycin (Sigma, P0781) at 37°C, 5% CO_2_, cells were used for mitochondria stress test using Seahorse Assay (Agilent, 103015-100). All isolation and culture steps were performed under the use of 10 µM contraction inhibitor (-)-blebbistatin (Targetmol, T6038) dissolved in DMSO.

For the Seahorse Assay, isolated cardiomyocytes were incubated with Seahorse XF DMEM medium, pH 7.4 (Agilent, 103575) supplemented with 0.68 mM L-glutamine (Gibco, 25030-081) and 5 mM glucose (Sigma G-7021) for 60 min at 37°C without CO_2_ prior to the Seahorse Assay. For substrate stimulation assays, seahorse medium contained either glucose and glutamine only, or additional 0.5 mM Na-butyrate (Sigma, 303410) or 1% (v/v) chemically defined (CD) lipids (including long-chain fatty acids; Gibco 11905031) or 0.1 mM MEM NEAA (Sigma, M7145) or 0.2 mM Na-3-hydroxybutyrate (BOH, Sigma, 54965). The Seahorse cartridge was prepared according to the manufacture protocol. Oxygen consumption rate (OCR) was measured for five cycles at baseline and after each injection (10 µM oligomycin [Sigma 75351], 10 µM FCCP [Sigma C2920], 1 µM rotenone/1µM antimycin A [Sigma R8875/A8674]) with settings of 1 min mix, 3 min wait, 3 min measure. For normalization to cell number, cardiomyocytes were washed with 1x PBS after the Seahorse Assay and stained using 1 µg/ml Hoechst (Sigma 33342), following measurement of fluorescence excitation and emission at 350 nm and 461 nm, using a plate reader Tecan Infinite Lumi 200 pro.

For isotope labeling stimulation with 0.5 mM ^13^C4 Na-butyrate (Sigma 488380), cells were incubated with previously described Seahorse XF DMEM medium, pH 7.4 for 90 min at 37°C, 5% CO_2._ After incubation, cells were washed with 0.9% NaCl, and frozen at −70°C until further mass spectrometric analysis.

### Cardiac ^13^C4 Na-butyrate isotope perfusion

For isotope labeling with ^13^C4 Na-butyrate (Sigma 488380), rat hearts from WT (n=3 ^12^C4 Na-butyrate WT; n=5 ^13^C4 Na-butyrate WT) and dTGR (n=5 ^12^C4 Na-butyrate dTGR) were perfused using the Langendorff-free isolation method^41^. A physiologic, Krebs-Henseleit-like solution (pH 7.4) was prepared containing 130 mM NaCl, 5 mM KCl, 0.5 mM NaH_2_PO_4_, 10 mM HEPES (Sigma, H3375-500), 10 mM Taurine, 1 mM MgCl_2_, 2 mM CaCl_2_, 1 % (v/v) CD lipids (Gibco 11905031, 100%), 1 % (v/v) insulin-transferrin-selenium (Sigma, I3146, 100x), 0.1 mM MEM NEAA (Sigma, M7145), 0.68 mM L-glutamine, 5 mM glucose, 0.1% bovine serum albumin (Sigma, A7906), 10% (v/v) FCS (PAN-Biotech, P30-3306), 0.1 mM Na-3-hydroxybutyrate (Sigma, 54965) and either 0.5 mM ^12^C4 Na-butyrate (Sigma, 303410) or 0.5 mM ^13^C4 Na-butyrate (Sigma 488380). In short, rats were decapitated after anesthesia with isoflurane. Thorax was opened and a cannula pierced into the apex to remove blood with 10 ml ^12^C4 Na-butyrate containing perfusion buffer while aorta and descending vena cava were cut open. Aorta and vein were closed above the atria with a clip and the heart was placed into a petri dish. The following steps were performed by injection into the LV: flushing with ^12^C4 Na-butyrate perfusion buffer (10 ml) to remove blood, 30 min perfusion with 20 ml recycling ^13^C4 Na-butyrate perfusion buffer, flushing with ^12^C4 butyrate perfusion buffer (10 ml) to remove excess ^13^C4 Na-butyrate from the heart. The heart was dissected into six LV regions as described in section Animal protocols, snap frozen and stored at −70°C.

### Gene expression analysis

Cardiac mRNA was isolated from the LV using Qiazol Lysis Reagent (Qiagen) and RNeasy Mini Kit (Qiagen) according to the manufacture protocol. For synthesis of cDNA from mRNA (2µg) the High-Capacity cDNA Reverse Transcription Kit (Thermo Fisher Scientific) was used according to manufactures protocol. Real-time PC was carried out with either TaqMan Fast Universal PCR or Fast Sybr Green (both Thermo Fisher Scientific) methodology and measured with QuantStudio 3 & 5 Real-Time PCR System (Applied Biosystems). The primer and probes were designed exon spanning with Primer Express 3.0 and ordered from BioTez GmbH. The relative standard curve method was used for quantification of target mRNA expression. Target gene expression was normalized to the expression levels of 36b4 housekeeping gene.

### Histology

Heart tissue was fixed in 4% buffered formalin solution for 48 hours. After dehydration with ethanol and xylol, organs were embedded in paraffin. The heart was cut into 2 µm thick slices with an electronic rotary microtome. Paraffin was removed before staining with xylol (3x 5 min) and a descending alcohol series (100 - 30% ethanol) was used for rehydration. Respective samples of one histological method were simultaneously stained and scanned with the Slide Scanner Panoramic MIDI immediately (objective plan-apochromat 20x/0.8x; Zeiss). Antigen unmasking was carried out by boiling samples (20 min) in citrate buffer, protein digestion was achieved using 3% hydrogen peroxide (15 min, RT) and 10% normal donkey serum (NDS) was used for blocking. Primary antibody (Rabbit anti-rat Fibronectin antibody [1:75] Ab23751; Abcam, Mouse anti-rat CD68 antibody | Clone ED1 [1:50] MCA341R; Bio-Rad) or biotinylated wheat germ agglutinin (WGA [1:100] B1025 VectorLab) or sirius red solution (0.1% in picric acid, Sigma) was diluted in 10% NDS and applied to the slice. After incubation in a humidity chamber overnight (4°C) or 60 min (Sirius red), excess antibody was removed, Cy3-conjugated secondary antibody (715-165-151, 711-165-152, respectively; Jackson ImmunoResearch) was diluted 1:100 or 1:300 in 1x PBS and incubated in a humidity chamber (120 min at RT). Excess antibody was removed and slides were mounted using Vectashield/Dapi medium. A negative control without secondary antibody was carried along.

### Proteomics

For label-free proteome analysis, LV and septal myocardial tissue was dissected in six regions: posterior base, mid, apex and anterior base, mid and apex. Tissue was snap frozen in liquid nitrogen and stored at −80°C. For peptide isolation approximately 30 mg tissue of each cardiac region were pulverized using using a CryoPrep CP02 (Covaris).

#### Sample Preparation and Protein Digestion

Peptide samples (100 µg per condition) were processed and digested using a modified single-pot, solid-phase-enhanced sample preparation (SP3) protocol^42^, automated on the AssayMap BRAVO system (Agilent Technologies). Protein lysates were reduced with 5 mM dithiothreitol (DTT) at 90°C for 10 min, alkylated with 10 mM iodoacetamide at room temperature for 30 min, and quenched with 20 mM DTT for 3 min. Paramagnetic beads (1 mg, Sera-Mag Speed Beads, Thermo Fisher Scientific) were added in 70% (v/v) acetonitrile, followed by an 18-min incubation and three washes with 70% ethanol. Proteins were digested overnight at 37°C using 4 µg of LysC and trypsin in 150 µl HEPES-KOH buffer (pH 7.6). Peptides were acidified with formic acid and desalted using the AssayMap Peptide Cleanup v2.0 protocol.

#### Liquid Chromatography–Mass Spectrometry (LC-MS) Analysis

Peptides (1 µg per replicate) were injected into an EASY-nLC 1200 system (Thermo Fisher Scientific) equipped with a 20 cm in-house packed analytical column containing C18-AQ 1.9 µm beads (Dr. Maisch Reprosil-Pur 120). Peptide separation was achieved using a 110-min reversed-phase gradient. Mass spectrometry analysis was performed on an Exploris 480 instrument (Thermo Fisher Scientific) operating in data-independent acquisition (DIA) mode with an asymmetric isolation window scheme.

#### Data Processing and Quantification

Label-free quantification was conducted using Spectronaut (Biognosys) version 16, implementing the directDIA workflow with default Pulsar search parameters. Spectra were matched against the UniProt rat proteome database (March 2022 release), including isoforms and common contaminants. Data from all heart segment samples were processed together. Protein quantification was performed using MaxLFQ normalisation^43^, and protein intensities were log₂-transformed.

Outlier replicates were excluded, and proteins with <75% valid values across all samples were filtered out. Remaining missing values were imputed using a randomized Gaussian distribution (width = 0.3, shift = 1.8), based on sample-specific means and standard deviations.

#### Statistical Analysis of Proteome

Differential abundance analysis was performed using the limma package^44^, applying empirical Bayes-moderated t-statistics. P-values were adjusted for multiple testing using the Benjamini-Hochberg method to control the false discovery rate.

Single-sample gene set enrichment analysis (ssGSEA)^45^ was performed to identify enriched pathways in the posterior apex of dTGR versus WT hearts (n = 4 dTGR, n = 5 WT), based on differential protein abundance. Log2 fold change (logFC) values were used as input, with protein groups collapsed to gene level by retaining the gene symbol associated with the lowest p-value. Duplicate gene symbols (n = 184) were resolved using this criterion. Gene sets were defined according to the Reactome database (m2.cp.reactome.v2024.1.Mm.symbols.gmt). ssGSEA was performed with rank-based sample normalisation, a weighting parameter of 0.75, and enrichment scoring based on the area under the running enrichment score curve (RES), using 10,000 permutations and a minimum gene set overlap of five genes. Normalised enrichment scores (NES) were reported, with statistical significance assessed by false discovery rate (FDR). Dot plots display the top 20 enriched Reactome pathways, with dot size proportional to gene set overlap (binned in 10% increments). Pathways were ranked by ascending nominal p-value, using descending absolute log₂ fold change (|logFC|) as a tie-breaker.

The visualization was done with ggplot2^46^, ComplexHeatmap^47^, Enhanced Volcano R packages (Blighe K, Rana S, Lewis M (2025). *EnhancedVolcano: Publication-ready volcano plots with enhanced colouring and labeling*. doi:10.18129/B9.bioc.EnhancedVolcano, R package version 1.27.0, https://bioconductor.org/packages/EnhancedVolcano).

### Metabolomics

#### Sample preparation

For metabolomic analysis, myocardial tissue regionally dissected and snap frozen as described before was pulverized using a BioPulverizer 59012MS (BioSpec), pre-cooled in liquid nitrogen prior every sample. Pulverized tissue was weighted and homogenized with zirconia beads in a Precellys 24 tissue homogenizer (Bertin). During homogenization and subsequent extraction, a mixture of methanol-chloroform-water (5:2:1 v/v/v), containing cinnamic acid as internal control, was used. Primary cardiomyocytes, myocardial biopsies, EHT and 2D iPSC cardiomyocytes were directly homogenized in methanol-chloroform-water. For metabolite extraction, cell lysates were shaken for 30 min at 4 °C, added water, shaken for another 30 min, and centrifuged for 15 min at 4 °C and 20800 x g. Polar, inter- and lipid phases were collected separately and dried under vacuum or nitrogen, respectively. Dried polar extracts were subjected to a second extraction in 20% (v/v) methanol and again dried under vacuum.

#### Untargeted metabolomics

For analysis of the central carbon metabolism, dried polar extracts were derivatized with 40 mg/mL methoxyamine hydrochloride in pyridine for 90 min at 30 °C and N-methyl-N-[trimethylsilyl]trifluoroacetamide (MSTFA) for 45 min at 37 °C. A mixture of nine alkanes (n-decane, n-dodecane, n-pentadecane, n-octadecane, n-nonadecane, n-docosane, n-octacosane, n-dotriacontane, and n-hexatriacontane) dissolved in hexane was added during derivatization to the MSTFA. A set of identification and quantification standards were derivatized and measured in parallel^48^. Measurements were performed on an autosampler (Gerstel)-coupled gas chromatography (Agilent 7890) time-of-flight mass spectrometer (Pegasus BT, Leco). The samples and identification/quantification standards were injected in split mode (split 1:10, injection volume 1 μL) in a temperature-controlled injector (CAS4, Gerstel) with a baffled glass liner (Restek). The following temperature program was applied during sample injection: initial temperature of 80 °C for 3 s, followed by a ramp with 7 °C/min up to 200 °C, a second ramp with 10 °C/min up to 250 °C, and final hold for 2 min. Gas chromatographic separation was performed on a Rxi-5 ms column of 30 m length, 250 μm inner diameter, and 0.25 μm film thickness (Restek). Helium was used as carrier gas with a constant flow rate of 1.2 ml/min. Gas chromatography was performed with the following temperature gradient: 2 min hold at 68 °C, first temperature gradient with 5 °C/min up to 120 °C, second gradient with 7 °C/min up to 200 °C, third gradient with 12 °C/min up to 335 °C, and a final hold of 5 min. Electron ionization was used with an electron energy of 70 eV. The transfer line temperature was set to 250 °C. Spectra were recorded for metabolites with 5 spectra per second at a detector voltage of 2060 to 2090 V and a mass range of m/z between 60 - 550 mu. Raw data was processed with ChromaTOF (version 5.55 BT). Peak integration was performed on the extracted ion chromatogram with a mass tolerance of 500 ppm and a minimum signal-to-noise of 10 and minimum stick count of 3. The peaks of the alkane mix were used for retention index calculation. An *in-house* library was used for spectral similarity library search on the identification standards with a minimum similarity score of 400, in order to generate a reference list. This reference list was then used for targeted annotation of metabolites in the quantification standards and samples. After annotation of metabolites in ChromaTOF, further analysis was performed using the *in-house* developed R-package MetaLabs (unpublished). In brief, quality of sample preparation and measurement was controlled based on internal standard abundance, sum of all areas per sample, derivatization mass abundance, alkane retention time alignment, metabolite annotation per sample, and quantification standards linearity. For visualization, the z-score per metabolite across all samples was calculated and depicted with ggplot2^46^ and ComplexHeatmap^47^.

#### Targeted metabolomics

For targeted analysis of the coenzyme A intermediates, dried polar extracts were resuspended in 5 mM hexylamine (pH=11.11). C18 ZipTip pipette tips (Merck Millipore) were used for desalting and concentration of samples. After washing and priming each ZipTip pipette tip with multiple methanol and 5 mM hexylamine strikes, samples were aspirated manifold times, ZipTip pipette tips washed with 5 mM hexylamine, nucleotides eluted in 0.3 mM ammonium acetate and 3.5 mM hexylamine in 27% methanol, and equal volumes of 100% methanol added to each sample. Quantification standards, containing a serial dilution of nucleotides and coenzyme intermediates in concentrations from 10 μM to 1 nM, were prepared and measured alongside the samples. Samples and standards were loaded onto a 96-well plate and run by direct infusion-tandem mass spectrometry, using a TSQ Quantiva Triple-Stage Quadrupole mass spectrometer (Thermo Fisher Scientific) coupled to a TriVersa NanoMate electrospray ionisation system (Advion). The ion spray voltage was set to 1.5 kV. Argon was used as a collision gas at a pressure of 1.5 mTorr and Q1 and Q3 were set with a FWHM resolution of 0.7. Data acquisition for each sample was performed over 3 minutes, with a cycling time of 3.3 seconds. Per sample, a total of 114 selected reaction monitoring scans were recorded, making up for 40 nucleotides (and nucleotide intermediates), 4 coenzyme A intermediates (plus different ^13^C-labelled variants), ribose 5-phosphate, and cinnamic acid as internal control. The measurement was carried out in negative mode, summing two transitions for each of the 46 metabolites. Data was evaluated with the Xcalibur Qual Browser software (Thermo Fisher Scientific) to verify quality of the measurements. Samples with low signal quality were omitted. Data was further extracted by an OpenMS package and analyzed using an *in-house* developed R-package (unpublished).

### Measurement of SCFA

SCFA were analyzed by gas chromatography-mass spectrometry (GC-MS). Each 5 mg of feces or 90 μl of serum or interstitual fluid were mixed with internal standard (100 μM crotonic acid (Sigma Aldrich)), 10 μl HCL (37 %, Sigma Aldrich) and 100 μL diethylether (Sigma Aldrich). Samples were shaken for 60 min at 1500 rpm and 25 °C and then centrifuged for 10 min at 1500 × g. 50 μl of the upper organic phase were transferred to a glass vial (Macherey-Nagel), containing 10μl N-tert-butyldimethylsilyl-N-methyltrifluoroacetamide (MTBSTFA, Sigma Aldrich). Samples were derivatized for 45 min at 80 °C and 600 rpm and subsequently incubated at room temperature for 24 hours. Serial dilutions of SCFA were processed and analysed in parallel for absolute quantification.

GC-MS analysis was performed on a Trace 1310 GC – Q Exactive MS, coupled to a TriPlus RSH autosampler (Thermo Fisher Scientific). Samples were injected in split mode (injection volume 1 μL for feces and 4 µL for serum or interstitial fluid, split 1:10). The following temperature program was applied during sample injection: initial temperature of 80 °C for 3 s followed by a ramp of 7 °C/sec to 210 °C and final hold for 3 min. Gas chromatographic separation was performed with a Rxi-5ms column (30 m length, 250 μm inner diameter, 0.25 μm film thickness (Restek)). Helium was used as carrier gas with a flow rate of 1.2 ml/min. Gas chromatographic separation was performed with the following temperature gradient: 2 min initial hold at 68 °C, first temperature gradient with 7 °C/min up to 150 °C, and second temperature gradient with 50 °C/min up to 300 °C, with a final hold for 2 min. The spectra were recorded in a mass range of 65 to 600 m/z with a resolution of 60,000. Data was analyzed with Xcalibur QuantBrowser software (Thermo Fisher Scientific), using serial dilutions of SCFA (ranging from 100 nM to 5 mM) as well as internal standard (crotonic acid) calibration for quantification (linear fit of area-ratio (SCFA/snternal standard) to concentration, with 1/X weighting). For butyrate mass 145.0680 m/z, 131.0524 m/z for propionate, and for crotonic acid mass 143.0523 m/z were monitored.

### Human UK Biobank cohort analysis

This analysis was conducted within the UK Biobank, a population-based cohort of 502,366 participants aged 40–69 years recruited between 2006 and 2010 across 22 assessment centres in England, Scotland, and Wales. Ethical approval was granted by the North West Multi-Centre Research Ethics Committee (reference 11/NW/0382), and all participants provided written informed consent. Data for this analysis were accessed on 16 November 2023.

Among all participants, 24,635 non-fasting individuals (blood sampling performed less than 5 hours after their last meal) fulfilled validated clinical criteria for heart failure with preserved ejection fraction (HFpEF) according to Versnjak et al.^32^, defined through a multi-stage algorithm conceptually aligned with the European Society of Cardiology (ESC) recommendations. Inclusion required at least one clinical symptom or sign of heart failure supported by objective evidence from cardiac magnetic resonance imaging, NT-proBNP concentration, or pre-test probability. When imaging data were unavailable, participants with NT-proBNP levels above the 90th percentile and a high pre-test probability of HFpEF (≥70%) were included, provided that heart failure with (mildly) reduced ejection fraction (HFmrEF and HFrEF) was absent. Underlying diagnostic information and censoring data were ascertained from linked hospital episode statistics, primary and secondary care records, and national death registries.

Arterial hypertension was defined using diagnostic codes mapped to the International Classification of Diseases, Tenth Revision (ICD-10), harmonized through the Observational Medical Outcomes Partnership (OMOP) Common Data Model. The definition followed clinical standards, relying on diagnostic labels rather than single blood pressure readings. Participants with a diagnosis of essential or secondary hypertension (ICD-10 I10–I15) recorded at or before baseline were classified as hypertensive, while all others were considered non-hypertensive. This approach minimized misclassification arising from blood pressure variability.

Baseline plasma samples were analyzed using quantitative nuclear magnetic resonance (NMR) spectroscopy (Nightingale Health, Finland). The platform quantified 251 metabolic biomarkers, including lipids, amino acids, and other metabolites related to energy metabolism. Circulating 3-hydroxybutyrate (BOH) was quantified in absolute molar units. Analytical reproducibility and quality control followed established Nightingale Health protocols (coefficients of variation <5%).

Analyses were stratified by sex to account for physiological differences in metabolic regulation and heart failure pathophysiology. Participants were further stratified by hypertension status, yielding four analytic subgroups: normotensive males, hypertensive males, normotensive females, and hypertensive females. Baseline characteristics were compared using linear regression for continuous variables and χ² tests for categorical variables.

The primary outcome was all-cause mortality, analyzed using Kaplan–Meier estimates and Cox proportional hazards models with the Breslow method for ties. The proportional hazards assumption was confirmed for all covariates. Hazard ratios (HRs) were reported with 95% confidence intervals (CIs). BOH concentrations were logarithmically transformed to approximate a normal distribution for continuous analyses and subsequently Z-standardised. Percentile-based groupings (<25th, 25th–75th, and >75th percentiles) were derived from untransformed concentrations and applied for group comparisons. Models were adjusted for age and body mass index within each subgroup. Statistical significance was set at p < 0.05.

### Statistical analysis

Statistical analysis was performed using GraphPad Prism (version 9.1.0) and R (version 4.4.1). Summary data is presented as mean ± SD if not indicated differently. The normality of data was tested with D’Agostino-Person or Shapiro-Wilk normality test. Outliers were identified with ROUT outlier test (Q=1%) and excluded from further analysis. Significance between two groups was tested using unpaired two-tailed Welch’s t-test, three groups were tested using more than two groups were tested with one-way ANOVA with Dunnett’s test for multiple comparisons (normally distributed data) or Kruskal-Wallis test with Dunn’s post-hoc test for multiple comparisons (non-normally distributed data). Two-way ANOVA with Dunnett’s or Sidak’s post-hoc test for multiple comparison was performed for analysis of three groups and two independent variables. The concrete data presentation format, independent biological replicates, number of experiments, statistical analysis including multiple comparison test and p-values are displayed in the respective Fig. legend. The p-values were reported as *P< 0.05, **P< 0.01, ***P< 0.001, ****P< 0.0001 and are depicted in the Figure.

## Data availability

Data available upon request

## Acknowledgement

S.M.K., N.B., P.F., A.M.C., F.E., G.G.S., S.K.F., M.K, N.W., M.G., P.M., R.D., S.S., D.N.M. were all supported by the Deutsche Forschungsgemeinschaft (DFG, German Research Foundation) – Project-ID 437531118 – SFB 1470. Several authors were supported by the Deutsches Zentrum für Herz-Kreislauf-Forschung (DZHK) - project numbers 81Z0100110, 81Z0100114, 81X2100189 for D.N.M., 81Z0100105, 81Z0100115 for N.H., 81Z0100204, 81Z0100228 for F.E., 81X3100210; 81X2100282 for G.G.S., 81Z0100113 for S.K.F., 81Z0100105, 81Z0100115, 81X2100190 for M.G., for 81Z0100111, 81X3100113 for S.S.. D.N.M. and N.W. were supported by the DZGIF (DZG Innovation Fund), Topic “Microbiome”. The German Ministry of Research, Technology and Space (BMFTR) supported members of the TAhRget consortium (project-ID: 01EJ2202D D.N.M., V.McP.) and (project-ID:01EJ2202A N.W. and H.B.). N.W. (#852796) and G.G.S. (ERC StG 101078307) are supported by the European Research Council under the European Union’s Horizon 2020 research and innovation program grant and N.W. by the Corona Foundation in the German Stifterverband. S.G. and S.K. were supported by the BMFTR funding Multimodal Clinical Mass Spectrometry to Target Treatment Resistance (MSTARS) and M.K. for VADYS-ME, Grant Number: 01EJ2406A.

## Acknowledgements

We thank Jutta Meisel, Gabriele N’diaye, Jana Czychi, Alina Deter, Martin Taube, Kerstin Puhl, Jenny Grobe and Mohamad Haji for assistance and technical support.

## Author contributions

S.M.K. and N.H. performed most experiments, and analyzed and interpreted the data. S.M.K., A.U., N.T. and N.H. performed the animal experiments, analyzed and interpreted the resulting data, with input from T.U.P.B.; S.Y.G. designed and performed the metabolomics and proteomics experiments and carried out data analysis and interpretation, with contributions from Y.D.Z., R.E-B., G.M. and G.N.K.; O.P. designed and performed further proteomics experiments and conducted proteomics data analysis and interpretation with support from A.F., N.B. and P.F.; A.U. performed gene-expression analyses and histology and analyzed and interpreted the data. S.M.K. analyzed the echocardiographic data with input from K.K. and L.P.; P.P. performed *in vitro* experiments and analyzed the data. M.K., A.M.C., C.T. and F.E. supported the translational approaches and provided critical input. S.M.K., S.Y.G., O.P., A.F. and N.H. performed the statistical analyses. P.M. and S.K. supervised the metabolomics and proteomics experiments and analyses and contributed to data interpretation. S.M.K., S.S., D.N.M., S.K. and N.H. conceived and designed the project, supervised the experiments, interpreted the data and acquired funding. S.M.K., S.Y.G., S.K., D.N.M. and N.H. wrote the manuscript, with editing by G.G.S., N.W., S.S., S.K.F., M.G. and R.D., and with further input from all authors.

## Extended data Figure

**Extended Data Fig. 1.**
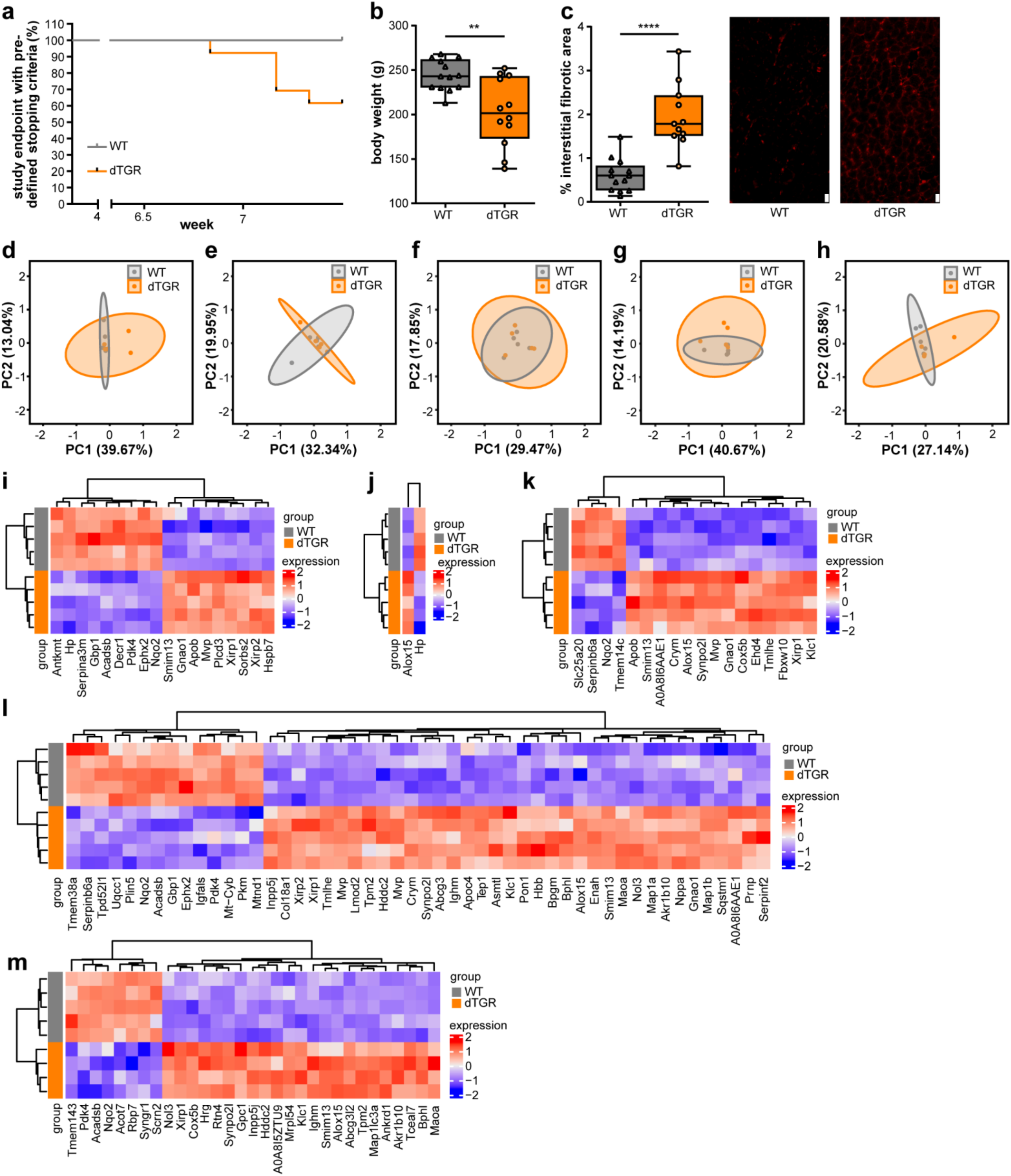
**(a)** Kaplan Meier curve of survival represents percentage of animals reaching pre-specified 3R termination criteria (n=13 WT, n=13-8 dTGR). **(b)** Body weight was significantly different between WT and dTGR (n=13 WT, n=12 dTGR). **(c)** Quantification of interstitial fibrotic area (left) (n=13 WT, n=11 dTGR) and representative images of immune histologic staining of fibronectin in myocardium (right; scalebar 20µm). Data are presented as boxplots (IQR) with whiskers min to max. *P<0.05, **P<0.01, ****P<0.0001 **(b-c)** unpaired two-tailed Welch’s t-test. **(d-h)** Principal component analysis (PCA) of all proteins with percentage of variance explained for different segments, **(i-m)** heatmaps displaying unsupervised hierarchical clustering between WT and dTGR of posterior base **(d, i)**, posterior mid **(e, j)**, anterior apex **(f, k)**, anterior mid **(g, l)**, anterior base **(h, m)**, adj.P<0.05, |logFC|>0.5, the expression data is scaled using z-scores for better visualization**. (d-m)** n=5 WT, n=4 dTGR

**Extended Data Fig. 2.**
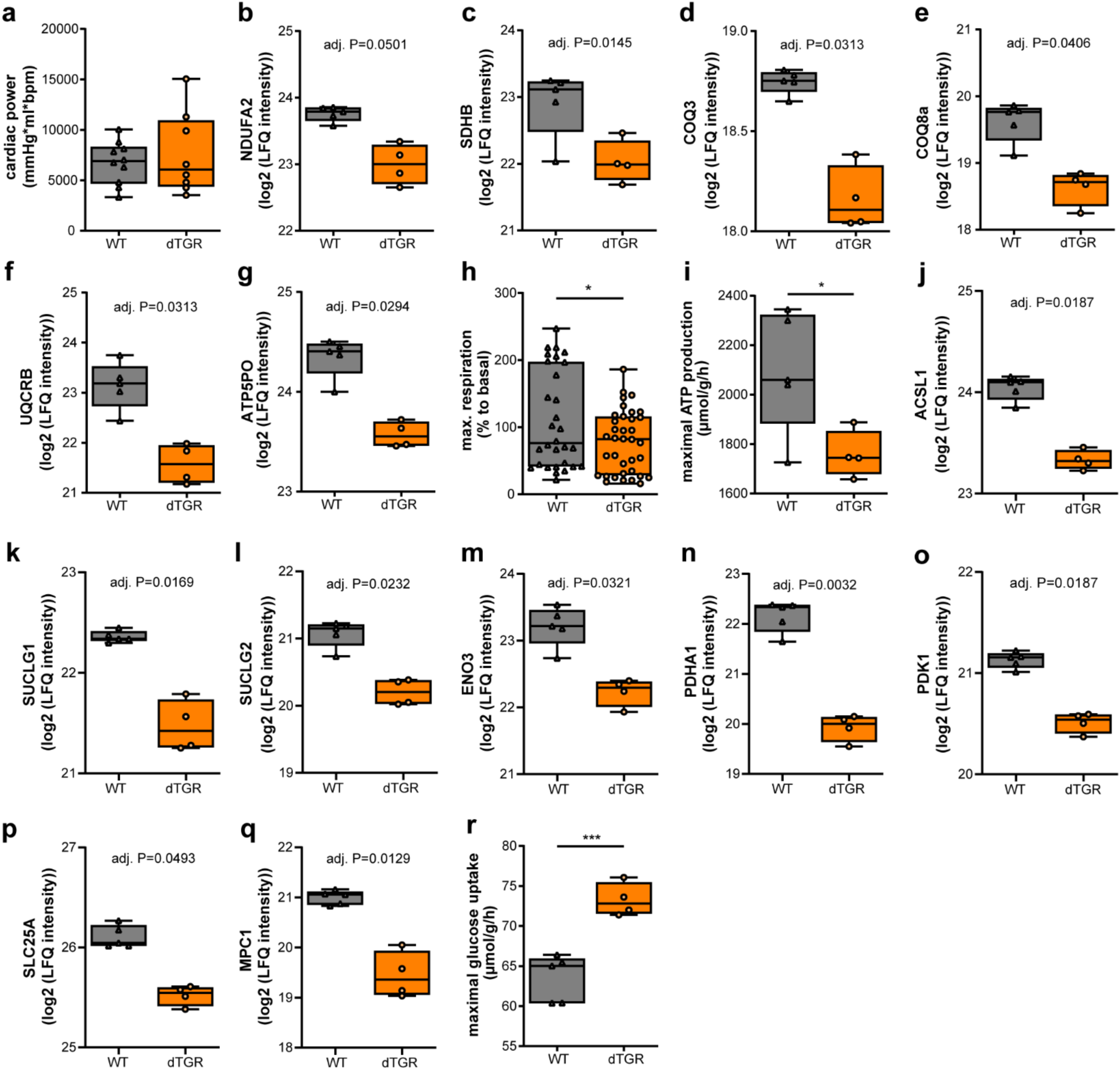
**(a**) Cardiac power (mmHg*ml*bpm) was similar in WT and dTGR hearts (n=10 WT, n=8 dTGR). **(b-g)** Boxplots depict protein intensities of proteins of the ETC with adjusted P-values from proteome analysis shown in the graphs (n=5 WT, n=4 dTGR). **(h)** Maximal respiration measured as pmol/min/Hoechst intensity with seahorse technology is displayed in percentage to basal respiration. (n=30 WT, n=36 dTGR technical replicates pooled from n=5-6 independent experiments **(i)** Maximal ATP production capacity and **(j)** maximal glucose uptake by metabolic modelling of metabolically relevant proteome using CARDIOKIN^26^. (n=5 WT, n=4 dTGR) **(k-r)** Boxplots depict protein intensities of proteins related to energy metabolism and substrate utilization with adjusted P-values from proteome analysis shown in the graphs. Data are presented as boxplots (IQR) with whiskers min to max. *P<0.05, ***P<0.001 unpaired two-tailed Welch’s t-test. **(h-i, r)**

**Extended Data Fig. 3.**
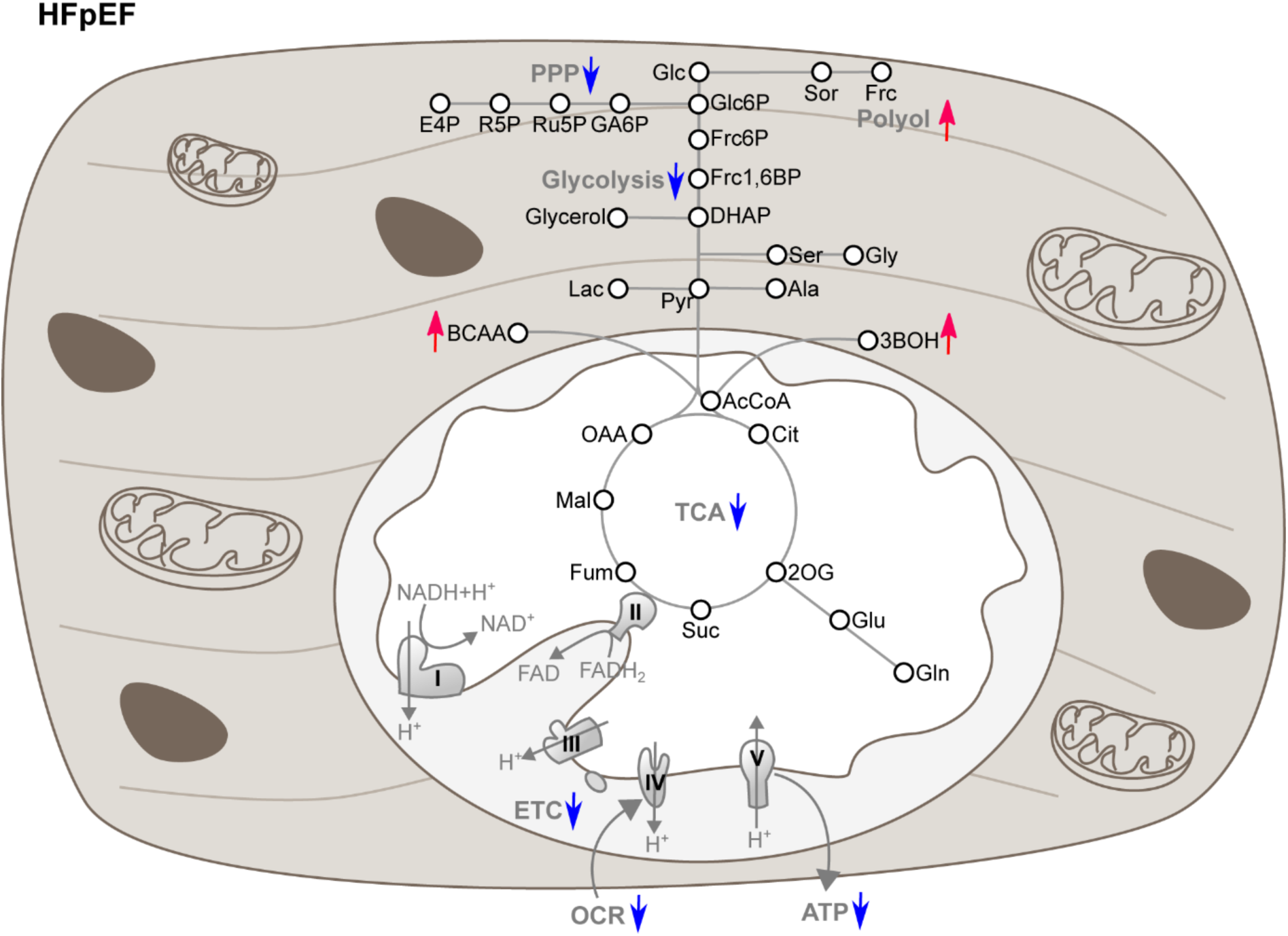
Pictogram of a HFpEF cardiomyocyte, with increased polyol intermediates, branched-chain amino acids and BOH (relative to WT, indicated as red arrows), while glycolysis, PPP, TCA cycle, ETC, OCR and ATP production are reduced (relative to WT, indicated as blue arrows)

**Extended Data Fig. 4.**
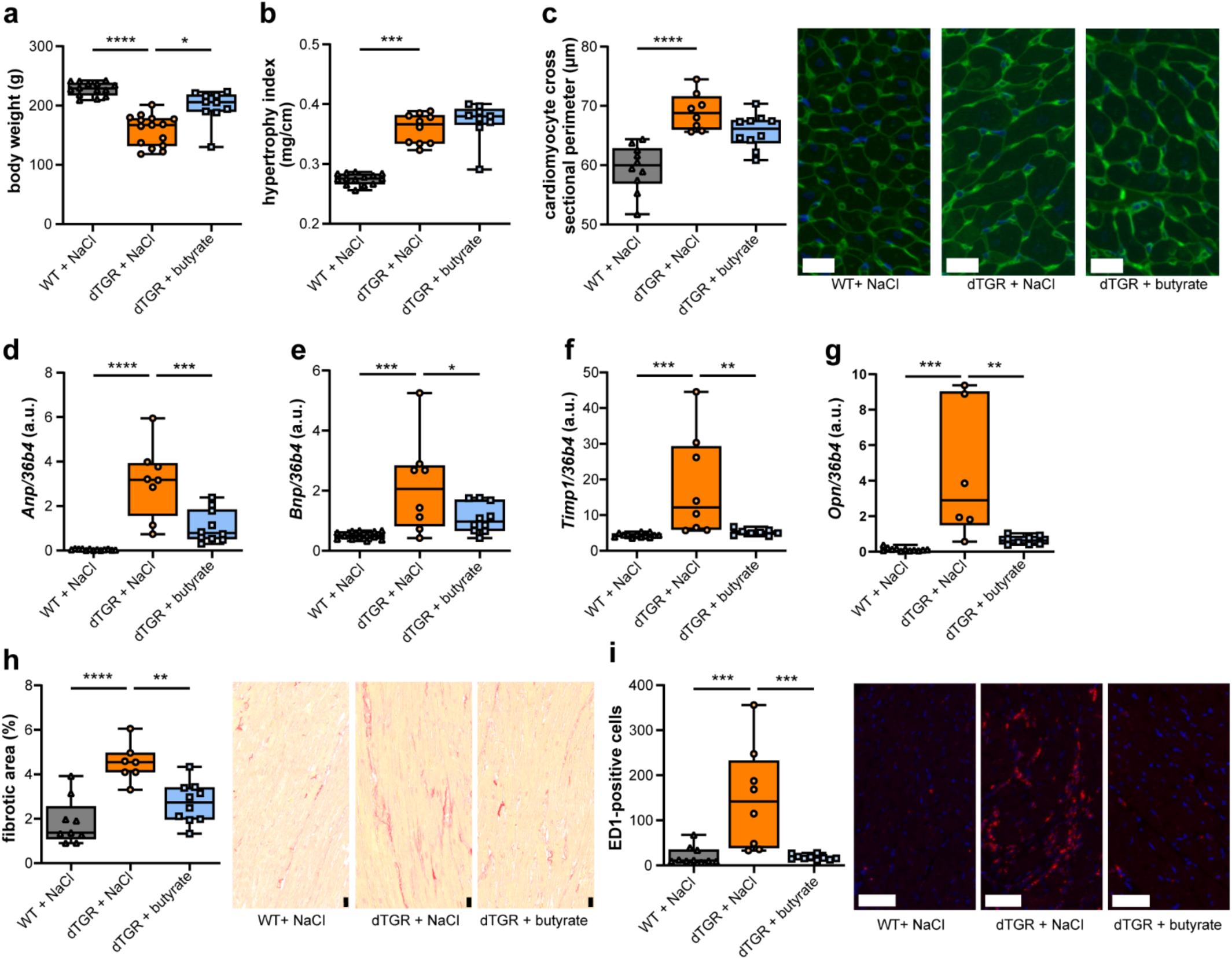
**(a)** Body weight was significantly different between WT and dTGR (n= 14 WT+NaCl, n=15 dTGR+NaCl, n=10 dTGR+butyrate). **(b)** cardiac hypertrophy index (heart weight to tibia length ratio) (n= 14 WT+NaCl, n=10 dTGR+NaCl, n=9 dTGR+butyrate. **(c)** Quantification of cardiomyocyte cross-sectional perimeter from WGA staining (left) (n= 10 WT+NaCl, n=8 dTGR+NaCl, n=10 dTGR+butyrate) and representative images (right, scalebar 20µm). Relative expression of **(d)** Anp, **(e)** Bnp, **f)** Timp, and **(g)** Opn to 36b4 measured by RT-qPCR (n= 13-12 WT+NaCl, n=8-6 dTGR+NaCl, n=10-8 dTGR+butyrate). **(h)** Quantification of fibrotic area from sirius red histologic staining (left) (n= 9 WT+NaCl, n=7 dTGR+NaCl, n=10 dTGR+butyrate), and representative images (right, scalebar 20µm). **(i)** Quantification of ED1-positive cells per field of view in ED1 (CD68) immune-histological staining (left) (n= 10 WT+NaCl, n=8 dTGR+NaCl, n=10 dTGR+butyrate), and representative images (right, scalebar 50µm. Outliers were removed upon statistical testing. Data are presented as boxplots (IQR) with whiskers min to max. **(a-i)**. *P<0.05, **P<0.01, ***P<0.001, ****P<0.0001 **(a)** Kruskal-Wallis test with Dunn’s multiple comparison, **(b-i)** ordinary one-way ANOVA with Dunnett’s multiple comparison.

**Extended Table 1:** ARRIVE 2.0 Compliance Checklist. This checklist summarizes compliance with the ARRIVE 2.0 guidelines for reporting animal research. The study described in the manuscript ’Butyrate Rescues Cardiac Metabolic Dysfunction in Hypertensive Heart Failure with Preserved Ejection Fraction’ followed these principles.

| ARRIVE 2.0 guidelines |
| --- |
| **Study design**<br>• Animals: WT (Sprague-Dawley) vs. double transgenic dTGR rats (SD-Tg(hREN)L10J × SD-Tg(hAGT)L1623)<br>• Groups:<br>◦ Cohort I untreated: WT, dTGR<br>◦ Cohort II treatment (200 mM NaCl or 200 mM Na-butyrate): WT+NaCl, dTGR+NaCl, dTGR+butyrate<br>• Module-specific subgroups: echocardiography, telemetry, Millar-tip<br>• Experimental unit: individual rat<br>**Section:** Methods: Animal protocols |
| **Sample size**<br>• Power calculations performed using G*Power<br>• $\alpha = 0.05$, Power = 0.80, SD and effect sizes predefined<br>• Example group sizes Cohort I:<br>◦ Echocardiography: WT n=13; dTGR n=13<br>◦ Millar-tip: WT n=10; dTGR n=9<br>• Example group sizes Cohort II:<br>◦ Echocardiography: WT+NaCl n=14, dTGR+NaCl n=16, dTGR+butyrate n=10<br>◦ Telemetry: dTGR+NaCl n=6; dTGR+butyrate n=5<br>◦ Millar-tip: WT+NaCl n=10; dTGR+NaCl n=9; dTGR+butyrate n=10<br>• Effect sizes (Cohen's d):<br>◦ Cardiac Output (CO): d = 1.19<br>◦ Blood pressure (BP): d = 2.32<br>◦ Arterial Elastance (Ea): d = 1.47<br>**Section:** Methods: Animal protocols, Figure legends |
| **Inclusion and exclusion criteria**<br>• Inclusion: male dTGR and WT controls, 4 weeks old, SPF except Pasteurella<br>• Exclusion: predefined welfare-related reasons, humane endpoints or unexpected clinical conditions unrelated to the experimental treatment<br>• Humane endpoints (summarized):<br>◦ A: mild signs (reduced general condition, pale mucosa) → 2× daily monitoring<br>◦ B: moderate signs (immobility, anorexia >6h, wound issues, pain) → treatment + 3× daily monitoring<br>◦ C: severe signs (apnoea, apathy, paralysis, Body Condition Score 1) → euthanasia within 1 h |
| ○ Moribund animals were euthanised immediately<br>• Outlier removal: ROUT Q=1%<br>**Section:** Methods: Statistical analysis |
| **Randomisation**<br>• Allocation by litter and body weight<br>• Confounding minimized by standardized environment and SOPs<br>**Section:** Methods: Animal protocols |
| **Blinding**<br>• Telemetry, non-invasive BP, echocardiography, Millar-tip, proteomics, metabolomics performed by blinded core facility staff where possible<br>• Gene expression and histology performed blinded where possible |
| **Outcome measures**<br>• Survival, blood pressure, echocardiography (EF, GLS)<br>• Hemodynamics (Millar-tip)<br>• Fibrosis, inflammation, molecular markers<br>**Section:** Results, Figures 1–5 |
| **Statistical methods**<br>• Normality: Shapiro–Wilk, D’Agostino<br>• Tests: Welch’s t-test, ANOVA (Dunnett), Kruskal–Wallis, ROUT<br>• Effect sizes: Cohen’s d<br>• Software: GraphPad Prism, R<br>**Section:** Methods: Statistical analysis, Figure legends |
| **Experimental animals**<br>• Species: Rattus norvegicus<br>• Strains: WT → Sprague-Dawley, dTGR → SD-Tg(hREN)L10J, SD-Tg(hAGT)L1623<br>• Age: 4–7 weeks<br>• Sex: males only (female phenotype unsuitable)<br>• Health status: SPF, Pasteurella-positive<br>• Source: MDC internal breeding<br>**Section:** Methods: Animal ethics and breeding |
| **Experimental procedures:**<br>• Telemetry implantation<br>• Metabolic cage urine collection<br>• Non-invasive blood pressure<br>• Echocardiography<br>• Millar-tip catheterisation<br>• Euthanasia<br>• Handling training<br>• Analgesia: Carprofen, Metamizol, local Lidocaine<br>**Section:** Methods: various subsections |
| **Results** |
| • All results reported with mean ± SD, p-values<br>• Effect sizes reported as above<br>**Section:** Results section; Figures 1–5, Figure legends |
| **Abstract**<br>• Summary of aims, model, treatment, major findings and mechanistic implications<br>**Section:** Summary |
| **Background**<br>• Rationale: HFpEF mechanisms, metabolic defects, microbiome-derived butyrate role.<br>**Section:** Summary, Main |
| **Objective**<br>• Mechanistic and functional testing of butyrate effects on cardiac energetics and diastolic dysfunction in HFpEF<br>**Section:** Summary, Main |
| **Ethical statement**<br>• Approved by LaGeSo Berlin (G0036/20)<br>• EU and German legislation followed<br>**Section:** Methods: Animal ethics |
| **Housing and husbandry**<br>• 22±2°C, 55±10% humidity, 12/12h light<br>• Enrichment: nesting material, tubes<br>• Group housing except telemetry (single-housed)<br>**Section:** Methods: Animal ethics |
| **Animal care and monitoring**<br>• Analgesia protocols (Carprofen, Metamizol, local Lidocaine)<br>• 2–3 days of handling training<br>• Daily monitoring + module-specific checks<br>• Humane endpoints applied as described<br>**Section:** Methods: Animal protocols |
| **Data access**<br>• Data available upon request |
| **Declaration of interests:**<br>• Funding and institutional support listed<br>• No competing interests declared<br>**Section:** Acknowledgements / Funding |

## References

1 Capone, F. et al. Cardiac metabolism in HFpEF: from fuel to signalling. Cardiovascular research 118, 3556–3575 (2023). 10.1093/cvr/cvac166

2 Neubauer, S. The failing heart--an engine out of fuel. The New England journal of medicine 356, 1140–1151 (2007). 10.1056/NEJMra063052

3 Murashige, D. et al. Comprehensive quantification of fuel use by the failing and nonfailing human heart. Science 370, 364–368 (2020). 10.1126/science.abc8861

4 Schonfeld, P. & Wojtczak, L. Short- and medium-chain fatty acids in energy metabolism: the cellular perspective. J Lipid Res 57, 943–954 (2016). 10.1194/jlr.R067629

5 Dinakis, E., O’Donnell, J. A. & Marques, F. Z. The gut-immune axis during hypertension and cardiovascular diseases. Acta physiologica, e14193 (2024). 10.1111/apha.14193

6 Boccella, N. et al. Transverse aortic constriction induces gut barrier alterations, microbiota remodeling and systemic inflammation. Sci Rep 11, 7404 (2021). 10.1038/s41598-021-86651-y

7 Santisteban, M. M. et al. Hypertension-Linked Pathophysiological Alterations in the Gut. Circ Res 120, 312–323 (2017). 10.1161/CIRCRESAHA.116.309006

8 Beale, A. L. et al. The Gut Microbiome of Heart Failure With Preserved Ejection Fraction. J Am Heart Assoc 10, e020654 (2021). 10.1161/JAHA.120.020654

9 Palm, C. L. et al. Beyond the gut: Systemic levels of short-chain fatty acids are altered in patients with heart failure. International journal of cardiology 428, 133124 (2025). 10.1016/j.ijcard.2025.133124

10 Jama, H. A. et al. Prebiotic intervention with HAMSAB in untreated essential hypertensive patients assessed in a phase II randomized trial. Nat Cardiovasc Res 2, 35–43 (2023). 10.1038/s44161-022-00197-4

11 Liu, P. et al. The role of short-chain fatty acids in intestinal barrier function, inflammation, oxidative stress, and colonic carcinogenesis. Pharmacol Res 165, 105420 (2021). 10.1016/j.phrs.2021.105420

12 Robles-Vera, I. et al. Protective Effects of Short-Chain Fatty Acids on Endothelial Dysfunction Induced by Angiotensin II. Front Physiol 11, 277 (2020). 10.3389/fphys.2020.00277

13 Panagia, M. et al. Increasing mitochondrial ATP synthesis with butyrate normalizes ADP and contractile function in metabolic heart disease. NMR Biomed 33, e4258 (2020). 10.1002/nbm.4258

14 Dong, T., Huang, D. & Jin, Z. Mechanism of sodium butyrate, a metabolite of gut microbiota, regulating cardiac fibroblast transdifferentiation via the NLRP3/Caspase-1 pyroptosis pathway. J Cardiothorac Surg 19, 208 (2024). 10.1186/s13019-024-02692-0

15 Yurista, S. R. et al. Ketone Ester Treatment Improves Cardiac Function and Reduces Pathologic Remodeling in Preclinical Models of Heart Failure. Circulation. Heart failure 14, e007684 (2021). 10.1161/CIRCHEARTFAILURE.120.007684

16 Carley, A. N. et al. Short-Chain Fatty Acids Outpace Ketone Oxidation in the Failing Heart. Circulation 143, 1797–1808 (2021). 10.1161/CIRCULATIONAHA.120.052671

17 Wilck, N. et al. Nitric oxide-sensitive guanylyl cyclase stimulation improves experimental heart failure with preserved ejection fraction. JCI Insight 3 (2018). 10.1172/jci.insight.96006

18 Ganten, D. et al. Species specificity of renin kinetics in transgenic rats harboring the human renin and angiotensinogen genes. Proceedings of the National Academy of Sciences of the United States of America 89, 7806–7810 (1992).

19 Luft, F. C. et al. Hypertension-induced end-organ damage : A new transgenic approach to an old problem. Hypertension 33, 212–218 (1999). 10.1161/01.hyp.33.1.212

20 Mervaala, E. M. et al. Endothelial dysfunction and xanthine oxidoreductase activity in rats with human renin and angiotensinogen genes. Hypertension 37, 414–418 (2001). 10.1161/01.hyp.37.2.414

21 Haase, N. et al. Relaxin does not improve Angiotensin II-induced target-organ damage. PLoS One 9, e93743 (2014). 10.1371/journal.pone.0093743

22 Wimmer, M. I. et al. Metformin modulates microbiota and improves blood pressure and cardiac remodeling in a rat model of hypertension. Acta Physiol (Oxf) 240, e14226 (2024). 10.1111/apha.14226

23 Heinzel, F. R., Hohendanner, F., Jin, G., Sedej, S. & Edelmann, F. Myocardial hypertrophy and its role in heart failure with preserved ejection fraction. J Appl Physiol (1985) 119, 1233–1242 (2015). 10.1152/japplphysiol.00374.2015

24 Tschope, C. & Senni, M. Usefulness and clinical relevance of left ventricular global longitudinal systolic strain in patients with heart failure with preserved ejection fraction. Heart failure reviews 25, 67–73 (2020). 10.1007/s10741-019-09853-7

25 Jani, V. P. et al. Myocardial Proteome in Human Heart Failure With Preserved Ejection Fraction. J Am Heart Assoc 14, e038945 (2025). 10.1161/JAHA.124.038945

26 Berndt, N. et al. CARDIOKIN1: Computational Assessment of Myocardial Metabolic Capability in Healthy Controls and Patients With Valve Diseases. Circulation 144, 1926–1939 (2021). 10.1161/CIRCULATIONAHA.121.055646

27 Tran, D. H. & Wang, Z. V. Glucose Metabolism in Cardiac Hypertrophy and Heart Failure. J Am Heart Assoc 8, e012673 (2019). 10.1161/JAHA.119.012673

28 Srivastava, S., Chandrasekar, B., Bhatnagar, A. & Prabhu, S. D. Lipid peroxidation-derived aldehydes and oxidative stress in the failing heart: role of aldose reductase. American journal of physiology. Heart and circulatory physiology 283, H2612–2619 (2002). 10.1152/ajpheart.00592.2002

29 Chen, S. et al. The role of glycolytic metabolic pathways in cardiovascular disease and potential therapeutic approaches. Basic research in cardiology 118, 48 (2023). 10.1007/s00395-023-01018-w

30 Jannapureddy, S., Sharma, M., Yepuri, G., Schmidt, A. M. & Ramasamy, R. Aldose Reductase: An Emerging Target for Development of Interventions for Diabetic Cardiovascular Complications. Front Endocrinol (Lausanne) 12, 636267 (2021). 10.3389/fendo.2021.636267

31 Solis-Herrera, C. et al. Effect of Hyperketonemia on Myocardial Function in Patients With Heart Failure and Type 2 Diabetes. Diabetes 74, 43–52 (2025). 10.2337/db24-0406

32 Versnjak, J. et al. Deep phenotyping of heart failure with preserved ejection fraction through multi-omics integration. European journal of heart failure (2025). 10.1002/ejhf.70041

33 Christensen, K. H. et al. Circulating 3-hydroxy butyrate predicts mortality in patients with chronic heart failure with reduced ejection fraction. ESC Heart Fail 11, 837–845 (2024). 10.1002/ehf2.14476

34 Krebs, H. A. The redox state of nicotinamide adenine dinucleotide in the cytoplasm and mitochondria of rat liver. Adv Enzyme Regul 5, 409–434 (1967). 10.1016/0065-2571(67)90029-5

35 Kraker, K. et al. Speckle Tracking Echocardiography: New Ways of Translational Approaches in Preeclampsia to Detect Cardiovascular Dysfunction. Int J Mol Sci 21 (2020). 10.3390/ijms21031162

36 Cerqueira, M. D. et al. Standardized myocardial segmentation and nomenclature for tomographic imaging of the heart. A statement for healthcare professionals from the Cardiac Imaging Committee of the Council on Clinical Cardiology of the American Heart Association. Circulation 105, 539–542 (2002). 10.1161/hc0402.102975

37 Klotz, S., Dickstein, M. L. & Burkhoff, D. A computational method of prediction of the end-diastolic pressure-volume relationship by single beat. Nature protocols 2, 2152–2158 (2007). 10.1038/nprot.2007.270

38 Takizawa, D. et al. Pathophysiologic and prognostic importance of cardiac power output reserve in heart failure with preserved ejection fraction. Eur Heart J Cardiovasc Imaging 25, 220–228 (2024). 10.1093/ehjci/jead242

39 Anand, V. et al. Prognostic value of peak stress cardiac power in patients with normal ejection fraction undergoing exercise stress echocardiography. Eur Heart J 42, 776–785 (2021). 10.1093/eurheartj/ehaa941

40 Avery, E. G. et al. Intestinal interstitial fluid isolation provides novel insight into the human host-microbiome interface. Cardiovascular research (2025). 10.1093/cvr/cvae267

41 Ackers-Johnson, M. & Foo, R. S. Langendorff-Free Isolation and Propagation of Adult Mouse Cardiomyocytes. Methods in molecular biology (Clifton, N.J.) 1940, 193–204 (2019). 10.1007/978-1-4939-9086-3_14

42 Hughes, C. S. et al. Ultrasensitive proteome analysis using paramagnetic bead technology. Mol Syst Biol 10, 757 (2014). 10.15252/msb.20145625

43 Cox, J. et al. Accurate proteome-wide label-free quantification by delayed normalization and maximal peptide ratio extraction, termed MaxLFQ. Molecular & cellular proteomics : MCP 13, 2513–2526 (2014). 10.1074/mcp.M113.031591

44 Ritchie, M. E. et al. limma powers differential expression analyses for RNA-sequencing and microarray studies. Nucleic Acids Res 43, e47 (2015). 10.1093/nar/gkv007

45 Subramanian, A. et al. Gene set enrichment analysis: a knowledge-based approach for interpreting genome-wide expression profiles. Proc Natl Acad Sci U S A 102, 15545–15550 (2005). 10.1073/pnas.0506580102

46 ggplot2 : Elegant Graphics for Data Analysis v. 2nd 2016. (Springer Springer International Publishing : Imprint: Springer, Cham, 2016).

47 Gu, Z., Eils, R. & Schlesner, M. Complex heatmaps reveal patterns and correlations in multidimensional genomic data. Bioinformatics 32, 2847–2849 (2016). 10.1093/bioinformatics/btw313

48 Opialla, T., Kempa, S. & Pietzke, M. Towards a More Reliable Identification of Isomeric Metabolites Using Pattern Guided Retention Validation. Metabolites 10 (2020). 10.3390/metabo10110457

